# Purifying Selection Shapes the Dynamics of P-element Invasion in *Drosophila simulans* Populations

**DOI:** 10.1101/2024.12.17.628872

**Authors:** Anna M. Langmüller, Benjamin C. Haller, Viola Nolte, Christian Schlötterer

## Abstract

**Background:** Transposable elements (TEs) are DNA sequences that can move within a host genome. Many new TE insertions have deleterious effects on their host and are therefore removed by purifying selection. The genomic distribution of TEs thus reflects a balance between new insertions and purifying selection. However, the inference of purifying selection against deleterious TE insertions from the patterns observed in natural populations is challenged by the confounding effects of demographic events, such as population bottlenecks and migration.

**Results:** We used Experimental Evolution to study the role of purifying selection during the invasion of the P-element, a highly invasive TE, in replicated *Drosophila simulans* populations under controlled laboratory conditions. Because the change in P-element copy number over time provides information about the transposition rate and the effect of purifying selection, we repeatedly sequenced the experimental populations to study the P-element invasion dynamics. Based on the empirical data, we used Gaussian Process surrogate models to efficiently explore the parameter space and identify parameter combinations that best reproduce the experimental P-element invasion trajectories. Assuming that beneficial P-element insertions are negligible, we found that, in our experimental populations, 73% (60.9% – 76.1%) of new P-element insertions are under purifying selection with a mean selection coefficient of −0.056 (−0.060 – −0.042), highlighting the central role of selection in shaping P-element invasion dynamics.

**Conclusion:** This study underscores the power of Experimental Evolution as a tool for studying transposable element invasions and highlights the pivotal role of purifying selection in regulating P-element dynamics.

## Background

Transposable Elements (TEs) are DNA sequences that can move and amplify within a host genome [1]. The evolutionary fate of TEs depends not only on their ability to replicate within the genome of their host, but also on the harm their activity might cause to their host [2]. Despite the challenges posed by their potentially disruptive behavior, TEs are ubiquitous in the tree of life [3,4], thriving in countless species, and have fascinated evolutionary biologists since their discovery more than seven decades ago [5].

It is widely accepted that some TEs insertions are harmful for their hosts [2,3,6], but the fraction of new TE insertions that are deleterious, and the distribution of their fitness effects, remain open questions [7,8]. Because measuring the effect of individual TE insertions on host fitness is tedious and a large number of independent insertions must be studied to obtain representative patterns, most evidence for purifying selection operating against TEs relies on the low abundance of TE insertions in specific regions of the host genome [9–11]. One limitation of this approach is the assumption of a uniform insertion probability of TEs across the genome. However, TEs can exhibit strong insertion preferences for specific genomic regions [3,12–14], which makes it difficult to distinguish between insertion bias and purifying selection.

An alternative method for detecting genomic signatures of purifying selection is based on the frequency spectrum of TEs within natural populations [15–17]. As with SNPs, demographic events such as population bottlenecks or admixture events pose a major challenge to the interpretation of TE frequency spectra [6,18]. Furthermore, recent TE activity can cause a shift in the site frequency spectrum that resembles the pattern expected from purifying selection, again making it difficult to distinguish between insertion bias and purifying selection [15,16,19]. Finally, the low population frequencies of TE insertions — caused by recent TE activity [16] and purifying selection — present a significant obstacle because detecting a specific TE insertion and estimating its frequency reliably requires high sequencing coverage.

Experimental Evolution (EE) [20,21], a powerful approach to study evolution under controlled laboratory conditions, is becoming increasingly popular as an alternative method to study TE invasion dynamics [22–24]. By combining EE with whole-genome sequencing of pooled individuals, an approach called Evolve and Resequence [25–27], researchers can minimize confounding factors that complicate the interpretation of polymorphism patterns in natural populations. Moreover, EE enables the replication of the invasion process, allowing stochastic (random) effects to be distinguished from deterministic patterns across multiple evolutionary replicates [20,21,26].

The P-element [28,29] — one of the best-studied TEs in eukaryotes [30] — is a 3 kb DNA transposon with a remarkable invasion record [31–33]. The P-element was most likely introduced into *Drosophila melanogaster* from *D. willistoni* through a horizontal transfer event and subsequently spread within *D. melanogaster* populations between the 1950s and 1980s [34]. It later spread to *D. melanogaster’s* sister species, *D. simulans*, via another horizontal transfer, spreading through *D. simulans* populations worldwide sometime between 2006 and 2014 [33,35]. The P-element consensus sequence in *D. simulans* and *D. melanogaster* differs by only a single base substitution at position 2040, where *D. simulans* has an ‘A’ and *D. melanogaster* has a ‘G’ [35]. This same polymorphism segregates in natural *D. melanogaster* populations, and recent analyses of P-element polymorphisms suggest that the horizontal transfer event from *D. melanogaster* to *D. simulans* likely occurred around Tasmania [36].

*Drosophila* species curb P-element invasions through the piRNA pathway, a specialized defense mechanism involving small RNAs that target and silence TEs [37,38]. Most piRNAs originate from specific genomic regions known as piRNA clusters, which collectively comprise approximately 3.5% of the *D. melanogaster* genome [37]. Current studies suggest that the effective silencing of P-elements across the genome relies on the presence of at least one P-element insertion within a piRNA cluster, which serves as a template for the production of P-element-specific piRNAs [39,40].

In this study, we used EE to investigate how purifying selection shapes the invasion dynamics of the P-element in experimental *D. simulans* populations. We followed the increase in P-element copy number in replicated EE studies across time. Using these time-resolved empirical invasion curves together with an individual-based simulation framework matching our experimental setup, we show that — under the assumption that P-element insertions are at most neutral and not beneficial [41] — only 27% (23.9% – 39.1%) of new P-element insertions can be considered as effectively neutral, while the remaining 73% (60.9% – 76.1%) are subject to purifying selection with an average selection coefficient of −0.056 (−0.060 – −0.042). The genome-wide mean selection coefficient for new P-element insertions outside piRNA clusters is −0.041 (−0.046 – - 0.030), highlighting the importance of purifying selection in shaping P-element invasion dynamics.

## Results

### The P-element rapidly invades experimental populations

To explore the influence of purifying selection on TE invasion dynamics, we analyzed the P-element invasion in replicate *D. simulans* populations from two EE experiments conducted under identical environmental conditions (Figure 1). These experimental populations were established from about 200 isofemale lines derived from a natural *D. simulans* population (Tallahassee, Florida). During the collection of the isofemale lines this natural *D. simulans* population was at the onset of a P-element invasion [35], where 25 – 44% of these isofemale lines were estimated to be P-element carriers [33]. The first EE experiment, which we refer to as the ‘1^st^ wave’ experiment, was started shortly after the isofemale line collection, thus all evolutionary replicates had only a small number of P-element copies at generation 0. The second EE experiment, which we refer to as the ‘2^nd^ wave’ experiment, on the other hand, was established about 4 years after the 1^st^ wave experiment from the same isofemale lines. During this period those isofemale lines that already contained a P-element at the time of collection, acquired additional P-element copies [11]. As a consequence, the 2^nd^ wave experiment started from a higher number of initial P-element copies. Because of the small effective population size during the maintenance of the isofemale lines, many of the P-element insertions that occurred in the isofemale lines between the setup of the 1^st^ and 2^nd^ wave experiment were deleterious. Purifying selection could only operate once the flies were maintained in large, outbred populations [11]. To track P-element invasion dynamics, we sequenced the replicate populations using the Pool-Seq approach [42] in both EE experiments and estimated P-element copy numbers per haploid genome using DeviaTE [43].

**Figure 1.**
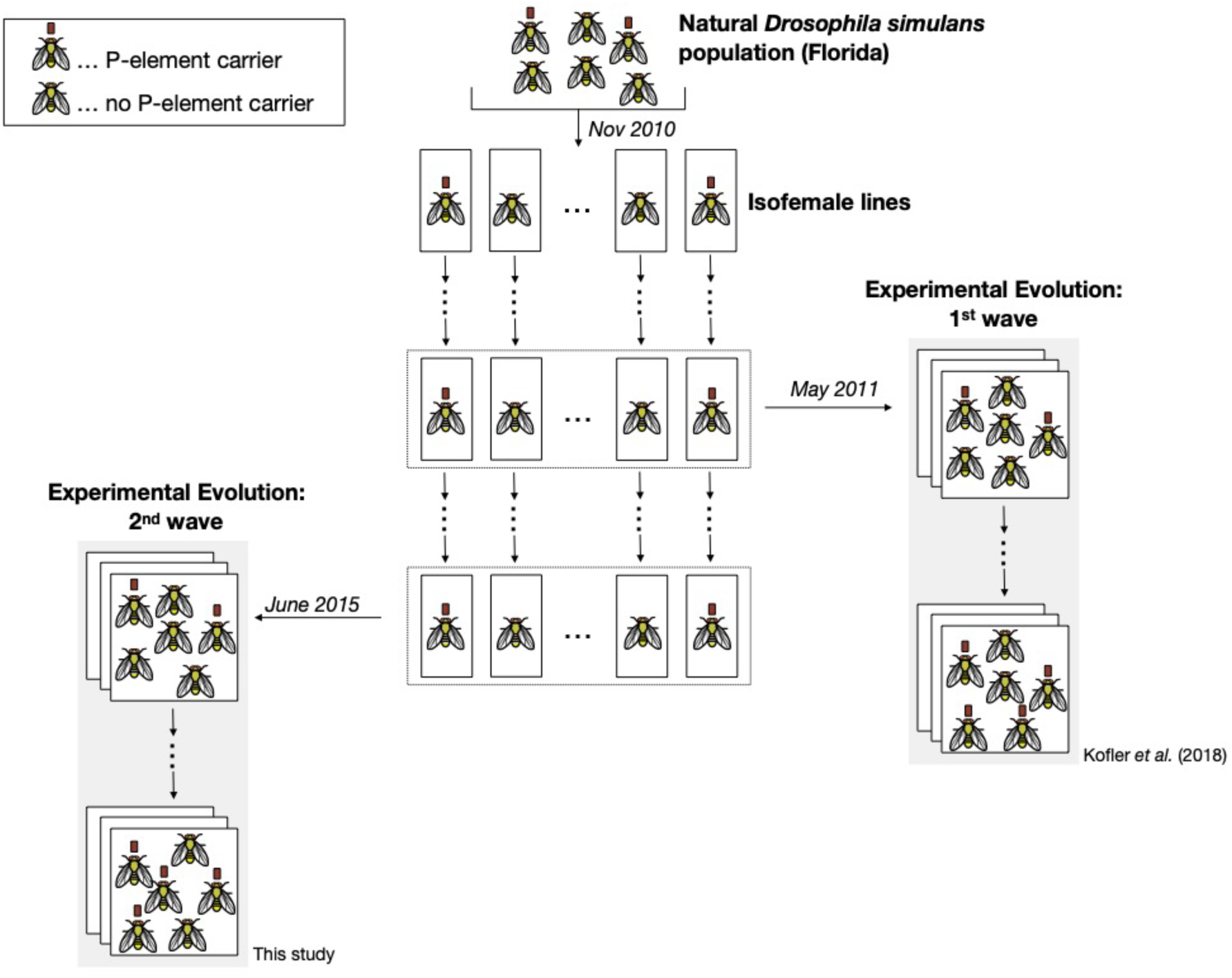
Schematic overview of the experimental design. Isofemale lines were established from a natural *Drosophila simulans* population, where the P-element has been invading [35]. Isofemale lines carrying an active P-element (indicated by red squares) are expected to accumulate P-element copies until an active defense mechanism is triggered [37,44]. In May 2011, three large replicate populations were established, using 202 isofemale lines. These populations, referred to as the “1^st^ wave” Experimental Evolution (EE) experiment, are maintained with non-overlapping generations under a cycling hot temperature regime and were sequenced every 10^th^ generation to monitor the P-element invasion dynamics [22]. In June 2015, a second EE experiment was initiated using three new replicate populations derived from 191 surviving isofemale lines [11]. These populations, referred to as the “2^nd^ wave” EE experiment, were maintained under the same environmental conditions and were also sequenced over time to track the P-element invasion.

Our data confirmed that the average initial P-element copy number in the 2^nd^ wave experiment was significantly higher than in the 1^st^ wave experiment (6.92 vs. 0.86, Figure 2). Despite distinct initial dynamics, both EE experiments reached a similar P-element copy number plateau: approximately 15 copies per haploid genome, after around 20 generations (Figure 2). This suggests that the timing and ultimate copy number plateau are consistent across both waves, indicating that these aspects of the P-element invasion are robust to initial conditions.

**Figure 2.**
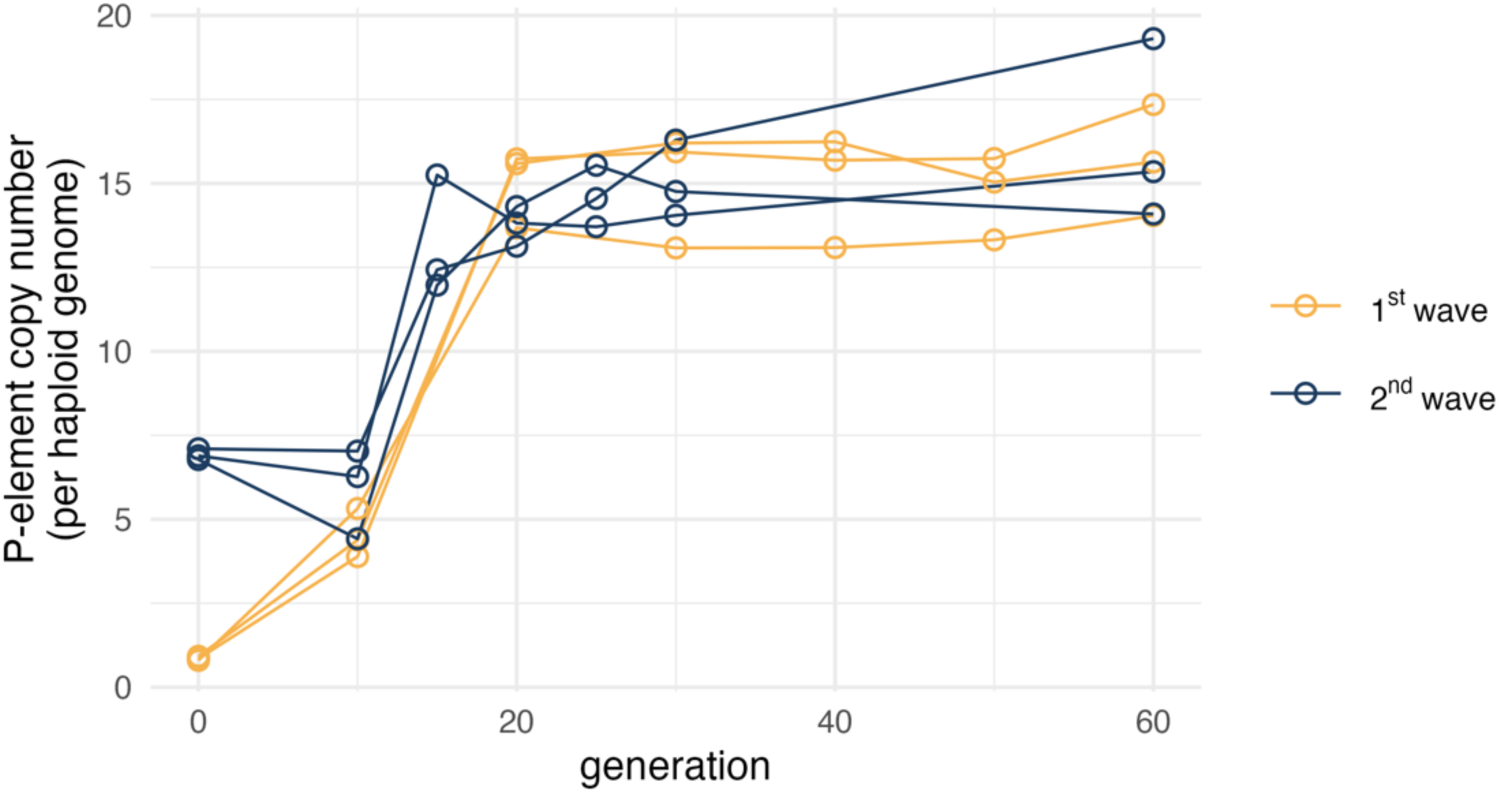
Estimated P-element copy number per haploid genome from the 1^st^ (amber) and 2^nd^ (dark navy blue) wave Experimental Evolution study. Sequenced time points are shown as circles.

### Purifying selection shapes P-element invasion dynamics

We estimated similar P-element copy numbers per haploid genome in generations 0 and 10 in the 2^nd^ wave experiment (means: 6.92 vs. 5.91, Figure 2). We previously hypothesized that this pattern arises from the balance between purifying selection, which removes deleterious P-element insertions, and transposition activity, which introduces new P-element copies [11]. Nevertheless, a comprehensive understanding of P-element invasion dynamics necessitates the availability of reliable estimates of the strength of selection operating on new P-element insertions. While genetic editing enables the inference of selection operating on specific TE insertions [45], it is evident that this approach is not feasible to describe the costs associated with insertions throughout the entire genome. Therefore, we used computer simulations to estimate the strength of purifying selection in our EE experiments.

#### Simulation framework

To investigate the potential influence of purifying selection on P-element invasion dynamics in the 1^st^ and 2^nd^ wave experiments, we developed an individual-based model for transposon dynamics in SLiM [46]. A comprehensive description of the model can be found in the Materials & Methods section. Figure 3 provides a schematic overview of the individual-based model, and Table 1 gives an overview of all model parameters. In brief, the individual-based model is able to simulate P-element invasions in EE experiments reflecting the empirical setups of the 1^st^ and 2^nd^ wave, respectively. As a host defense mechanism, we implemented a classic “trap model” [39,40,44]: P-elements transpose with a defined probability per copy and generation unless silenced by an insertion in a piRNA cluster [37], which immediately inactivates all P-elements in the genome.

**Figure 3.**
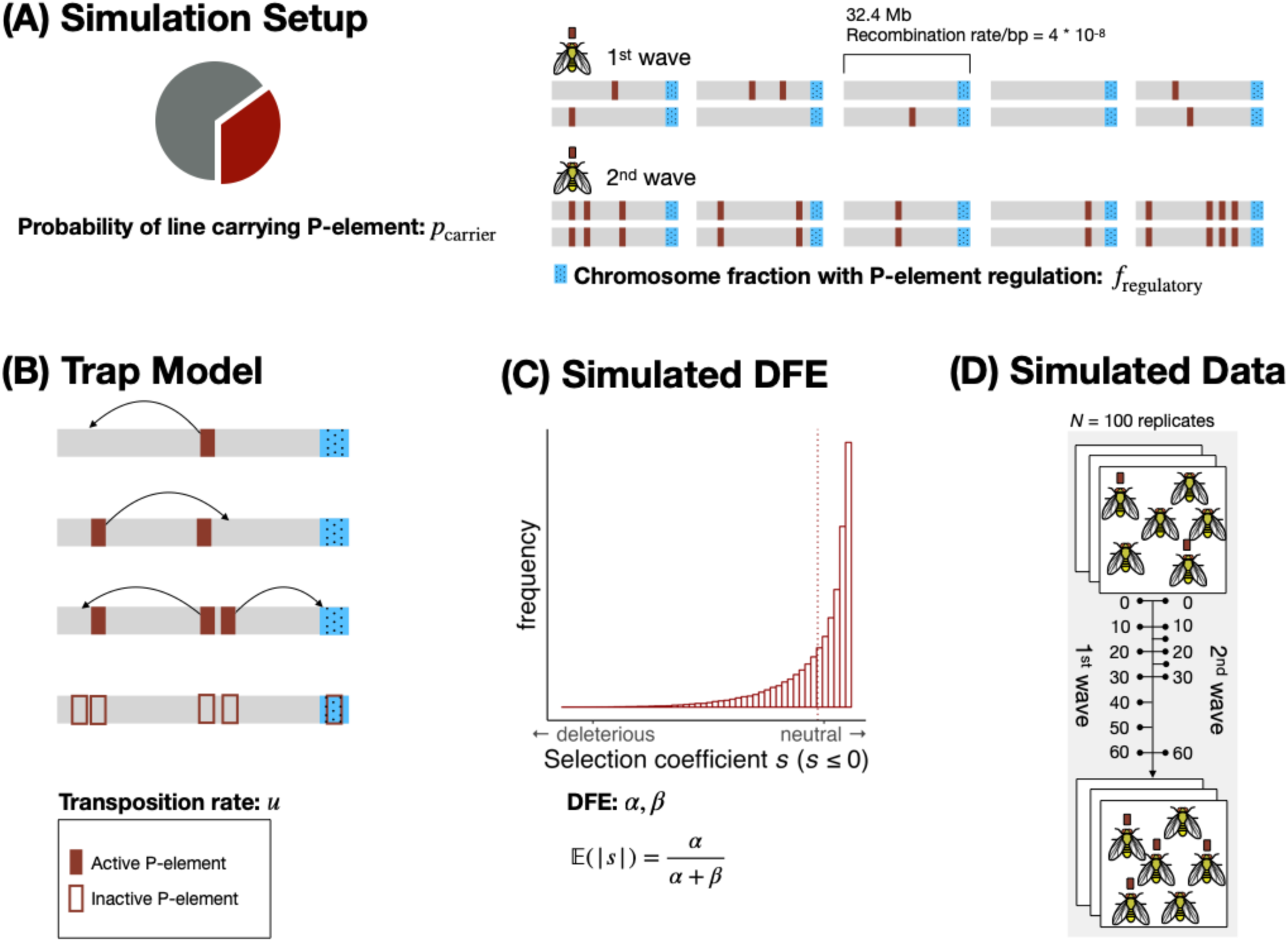
Schematic overview of the individual-based simulation model mimicking the P-element invasion dynamics in our Experimental Evolution experiments. **(A) Simulation Setup:** Ancestral outbred populations are generated by mixing 200 isofemale lines (five flies each). Each line carries the P-element with probability *p*_carrier_. We modeled diploid individuals with five chromosomes, each with a fixed length of 32.4 Mb and a recombination rate of 4×10⁻⁸ per bp per generation. For the 1^st^ wave, P-element insertions are assumed to be heterozygous. For the 2^nd^ wave, since the experiment started with isofemale lines that had been maintained at small populations sizes for 4.5 years [11], the model assumes that all P-element insertions are homozygous due to increased inbreeding and the likely establishment of a defense mechanism. The parameter *f*_regulatory_ defines the fraction of each chromosome with P-element-regulatory properties (piRNA clusters; blue rectangles). **(B) Trap Model:** The P-element remains active (filled rectangle) unless one of its copies transposes into a piRNA cluster (blue rectangles). The probability that a single P-element undergoes transposition in a given generation is controlled by the transposition rate *u*. Once a piRNA cluster acquires a single P-element insertion, all P-elements in the genome are immediately inactivated (unfilled rectangles with red borders). **(C) Simulated Distribution of Fitness Effects (DFE):** The DFE for new P-element insertions is modeled using a beta distribution with shape parameters *α* and *β*, which define the selection coefficient *s*. Positive selection is not considered in our model. **(D) Simulated Data:** The simulation output used in the analyses contains the average P-element copy number per haploid genome across 100 simulation runs, taken at the same time points used in the two Experimental Evolution studies. For a schematic of the extended model including parameters for dominance (*h*) and excision probability (v), see Supplementary Figure S1.

**Table 1.** Model parameters of the individual-based P-element invasion model and their considered ranges. The range for *p*_carrier_ was guided by a previous study estimating that 25 – 44% of the isofemale lines used in the 1^st^ and 2^nd^ wave experiment carried the P-element [33]. The range for *f*_regulatory_ was based on the estimate that 3.5% of the *Drosophila melanogaster* genome consists of piRNA clusters with TE-regulatory properties [37]. However, because piRNA clusters are challenging to assemble and compare across species [49] and because a previous simulation study suggests that as little as 0.2% may be sufficient for TE control [50], we allowed *f*_regulatory_ to vary rather than fixing it. The range for *u* was informed by the observed P-element invasion dynamics in the 1^st^ wave experiment. Specifically, the lower bound of 0.15 was chosen based on effective transposition rate estimates derived from generations 0 and 10 in the 1^st^ wave experiment. The upper bound of 0.5 ensures that effective transposition remains possible even under strong purifying selection scenarios (please refer to the Materials C Methods section for more detail). See Supplementary Table S1 for an overview of model parameters in the extended model, including dominance (*h*) and excision probability (*v*).

| Parameter | Description | Range |
| --- | --- | --- |
| $p_{\text{carrier}}$ | The probability that an isofemale line carries the P-element at the beginning of the simulation | [0.15, 0.5] |
| $f_{\text{regulatory}}$ | piRNA cluster size: the fraction of the chromosome with P-element-regulatory properties (once at least one P-element insertion is acquired) | [0.01, 0.1] |
| $u$ | The probability of transposition of a single P-element per generation | [0.15, 0.5] |
| $\alpha$ | The shape parameter $\alpha$ of the beta distribution that defines the selection coefficient $s$ for P-element insertions outside piRNA clusters | [0.001, 0.5] |
| $\beta$ | The shape parameter $\beta$ of the beta distribution that defines the selection coefficient $s$ for P-element insertions outside piRNA clusters | [10, 20] |

Outside piRNA clusters, P-element insertions are subject to purifying selection. For the main analysis, we assumed co-dominance by fixing the dominance coefficient at *h* = 0.5. We sampled the selection coefficient *s* for each P-element insertion from a beta distribution and multiplied *s* by −1 to ensure negative values (*s* ≤ 0). Wildtype homozygotes have a fitness of 1, individuals with a single heterozygous P-element insertion have fitness 1 + *hs*, and homozygotes for the P-element insertion have fitness 1 + *s*. By modifying the parameters of the beta distribution, it is possible to create a gradual transition between simulation scenarios where the majority of new P-element insertions are effectively neutral and scenarios where the majority of P-elements are subject to strong purifying selection, assuming no beneficial P-element insertions [41].

To explore the robustness of our result, we also implemented an extended version of the individual-based model that includes two additional free parameters: the dominance coefficient (*h*) and the probability of a clean excision event at the original insertion site, given that a P-element transposes (*v*). See Figure S1 for a schematic overview of the extended model, and Table S1 for an overview of the corresponding parameters. Results from this extended model are presented in the Supplement.

By comparing simulated invasion outcomes to empirical data and quantifying the fit, systematic exploration of this model’s parameter space (Table 1) would in principle allow us to determine the level of purifying selection that best fits the observed P-element copy number trajectories from the 1^st^ and 2^nd^ wave experiments. However, exhaustively probing our five-dimensional parameter space in this manner would be very computationally intensive [47], even using an optimized model. To address this challenge, we used Gaussian Process (GP) surrogate modeling [48].

A GP is a statistical model that serves as a surrogate for the individual-based model, allowing rapid prediction of the individual-based model’s behavior based on previously simulated data. The utility of the GP approach lies in its capacity to extrapolate from sparse data, thereby predicting the behavior of a system, such as our individual-based model, at untested parameter combinations. We trained two separate GPs — one for the 1^st^ wave and one for the 2^nd^ wave — using simulation outcomes from the individual-based model as described further in the Materials C Methods section. We then explored the accuracy of our trained GPs with a test dataset of 5000 additional simulation outcomes spanning a broad range of parameter combinations, for both the 1^st^ and 2^nd^ wave experiments. For this test dataset, the GPs accurately predicted trajectories of P-element copy numbers across generations (Figure 4, extended model: Figure S2). Based on these results, we conclude that the GPs provide an efficient and precise alternative to the individual-based model. This allowed us to use the GPs to find the model parameters that best replicate the experimental data, ultimately allowing us to estimate the level of purifying selection for which our individual-based model best fits the empirical time-series data.

**Figure 4.**
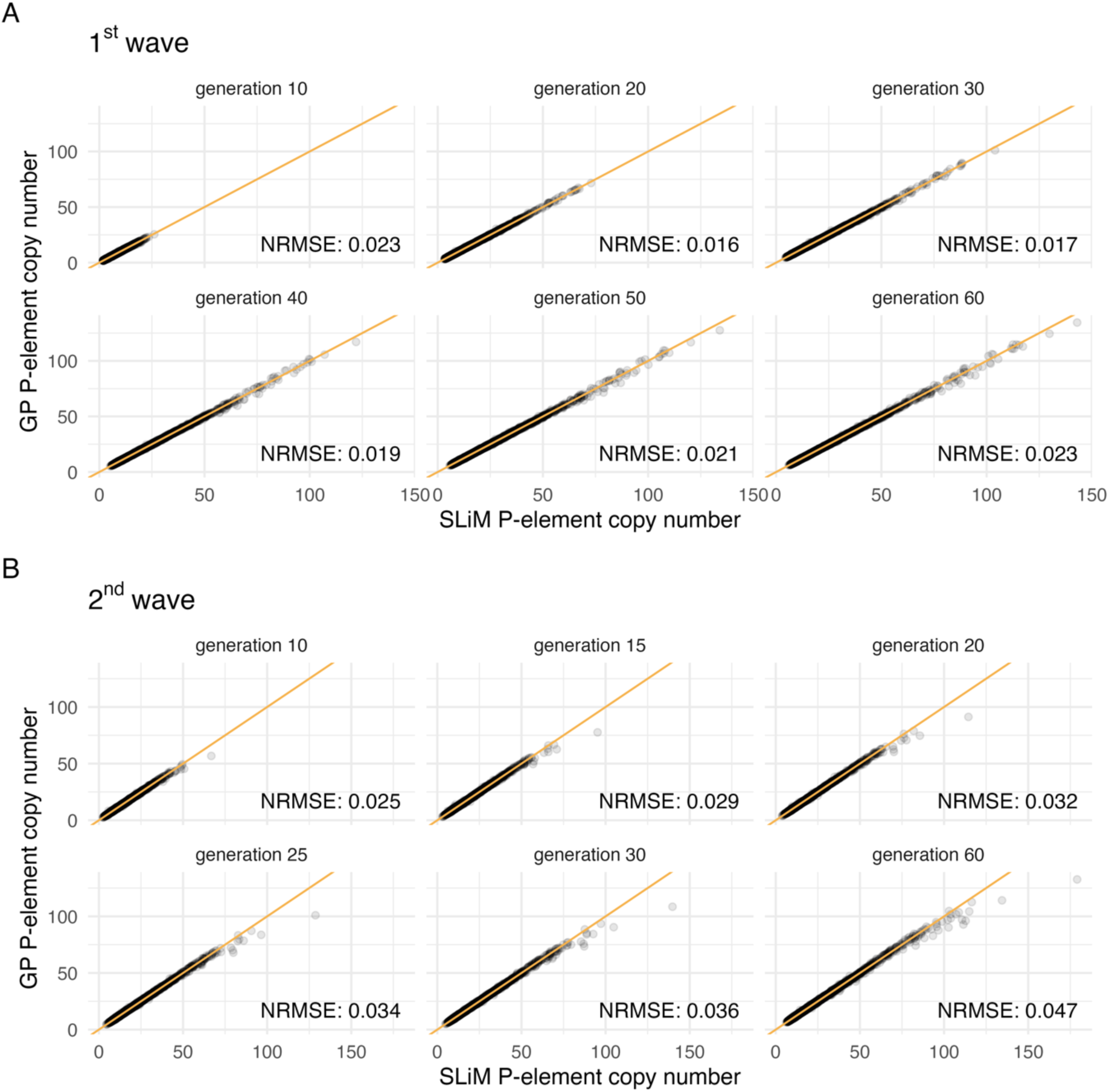
Gaussian Process (GP) performance: A test dataset consisting of 5000 data points was simulated with the individual-based model (*x* axis) for the (A) 1^st^ wave and (B) 2^nd^ wave and compared to the predictions of the GP (*y* axis). One data point in this test dataset comprises a specific combination of five model parameters (Table 1) and the corresponding predicted P-element copy numbers for six time points. The observed and predicted P-element copy numbers (gray dots) are shown for each of those six time points (the six panels), with amber lines indicating the identity line (*x = y*). Normalized root mean square errors (NRMSE) for the time points are shown at the bottom right of the corresponding panels. The analysis shows that the GP can predict the copy number observed in the individual-based model very accurately. Only at extremely high copy numbers — well beyond the empirical estimates (Figure 2) — can a slight underestimation be observed. Results for the extended model version including parameters for dominance (*h*) and excision probability (*v*) are shown in Supplementary Figure S2.

#### Purifying selection as a central mechanism: model calibration results

After validating the performance of the GPs with the simulated test dataset, we used the GPs to predict P-element invasion dynamics — the P-element copy number per haploid genome for six evolved time points corresponding to the sequenced time points of the empirical EE experiments — across 10^6^ model parameter combinations.

For each EE experiment (1^st^ and 2^nd^ wave), we generated predictions separately and compared them to the empirical data. Specifically, at each evolved time point, we compared the GP-predicted P-element copy number to the observed P-element copy numbers from all three biological replicates. At each evolved time point, we calculated the root mean square error (RMSE) between the predicted (n=1) and observed (n=3) values across replicates and normalized it by the mean observed copy number at that time point. These normalized RMSE (NRMSE) values — each reflecting the deviation in P-element copy number between the GP prediction and the three biological replicates at a single generation — were then summed across all time points to yield an overall prediction error score for each parameter combination. This approach ensures that later generations, which tend to have higher copy numbers (Figure 2), did not disproportionately influence the fit metric. Lower NRMSE values indicate a better agreement between the empirical and predicted P-element invasion curves for a specific parameter combination. We refer to the parameter combinations with the lowest NRMSE values as “top-ranked” parameter sets, meaning those that yielded the best agreement between GP predictions and empirical data within the predefined set of evaluated combinations.

We acknowledge that our approach identifies the parameter combination resulting in the best agreement between GP predictions and experimental data within the evaluated set, and that unexplored parameter combinations could, in principle, result in an even better agreement. Additionally, small differences in NRMSE among parameter combinations with low NRMSE values may not be indicative of substantively different predicted invasion dynamics and should not be overinterpreted. Thus, our NRMSE-based approach is a heuristic for ranking parameter plausibility rather than inference.

By comparing GP predictions from 10^6^ parameter combinations to the 1^st^ wave invasion dynamics, we observed a wide range of NRMSE values, spanning from 0.60 to 39.28. The best agreement, characterized by the lowest NRMSE, was observed for a parameter combination with notable purifying selection against new P-element insertions outside piRNA clusters (*s̅* = −0.045; range across the top 100 parameter combinations ranked by lowest NRMSE: (−0.047 – −0.036), Figure 5). Although the initial invasion dynamics of the 1^st^ and 2^nd^ wave EE experiments differ considerably, we reasoned that the genome-wide distribution of fitness effects for P-elements should be constant between the two experiments. Thus, we used the top-ranked parameter combination from the 1^st^ wave experiment to predict the invasion trajectory of the 2^nd^ wave experiment. Interestingly, this resulted in a remarkably accurate prediction of the invasion dynamics of the 2^nd^ wave experiment, which has been conducted four years later with an 8-fold higher initial P-element copy number (Figure 5). This result highlights the robustness and predictive power of our top-ranked parameter combinations even for rather distinct invasion trajectories in the first generations. Since the 1^st^ wave simulations lack homozygous P-element insertions at generation 0 and homozygous sites are unlikely to arise later due to transposition and outbreeding, we did not attempt to calculate top-ranked parameter combinations for the extended model (Figure S1) from the 1^st^ wave data alone, as it provides insufficient information to estimate the dominance coefficient *h*.

**Figure 5.**
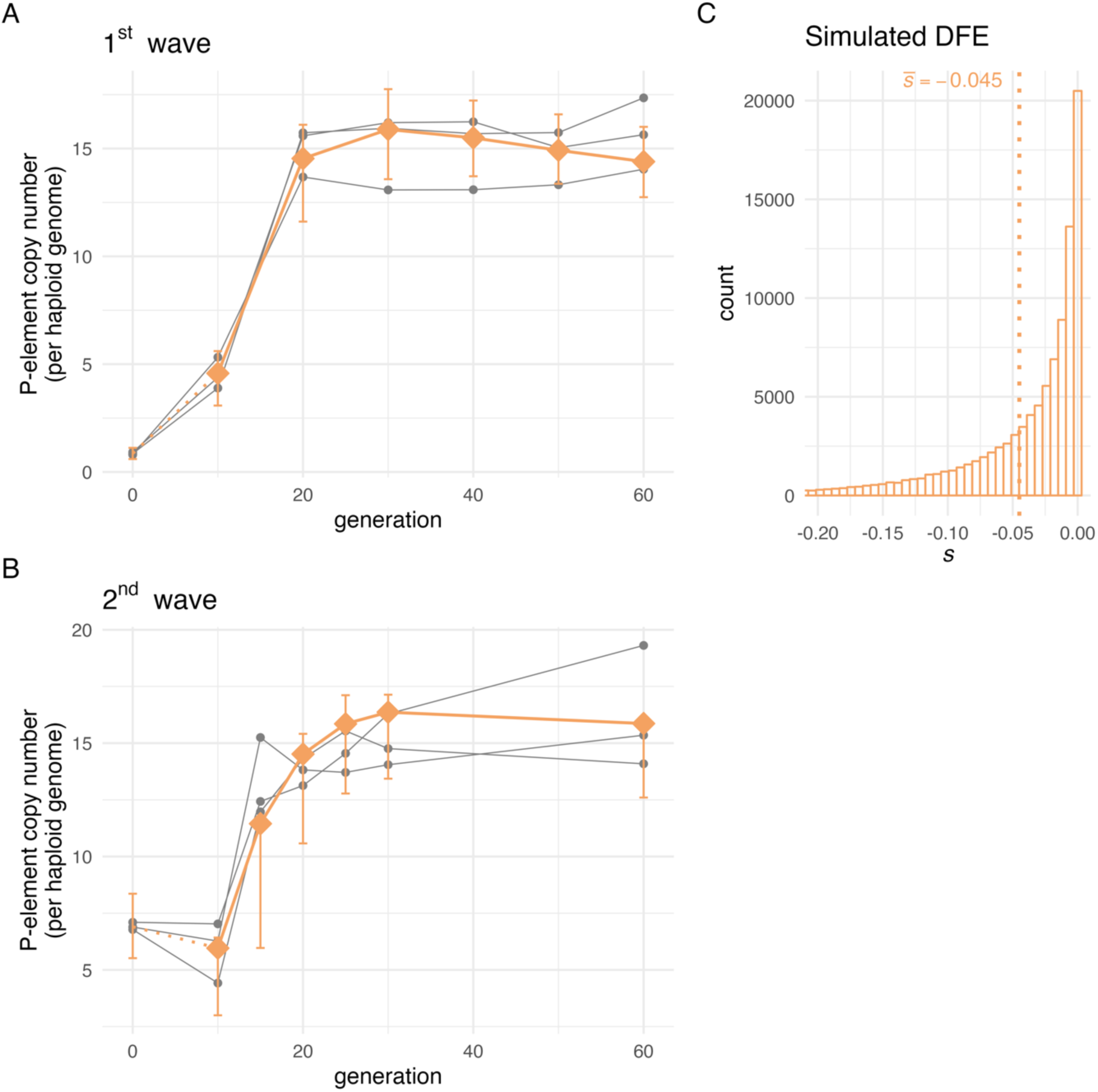
Gaussian Process (GP) predictions provide a good fit to the empirical data — the top-ranked parameters were identified using only the 1^st^ wave data: (A) 1^st^ wave experiment (B) 2^nd^ wave experiment. Each grey line represents an empirical evolution replicate, with sequenced time points indicated by dots. GP predictions are indicated by connected amber diamonds. The amber error bars show the range between the 2.5^th^ and 97.5^th^ percentiles of P-element copy number trajectories simulated with the individual-based model. These simulations were run with the same parameters as those used for GP prediction. The error bars are shown at the same time points where empirical data were sequenced. Note that GP predictions do not cover generation 0; instead, predictions from generation 10 are connected to the individual-based simulation averages at generation 0 to aid visual interpretation. (C) The simulated distribution of fitness effects (DFE) used for generating predictions in panels (A) and (B). Due to selection, the actual distribution of selection coefficients for segregating P-element insertions in the simulated populations will be skewed toward 0. The parameter combination used for the GP prediction is: *p*_carrier_ *= 0.18S, f*_regulatory_ *= 0.014, u = 0.277,* α *= 0.484,* β *= 10.327.* Predictions based on parameters identified from the 1^st^ wave accurately describe the P-element invasion dynamics in the 2^nd^ wave.

After confirming that the top-ranked parameters from the 1^st^ wave also explain the 2^nd^ wave data, we proceeded to jointly calibrate the model using both EE experiments. We reasoned that this could further refine our approach because our fit metric will be based on twice the amount of data. The overall fit metric was calculated as the sum of the NRMSEs for the 1^st^ and 2^nd^ wave experiments (NRMSE_sum_). We observed a considerable degree of variation in NRMSE_sum_ across the evaluated parameter combinations, with values ranging from 1.340 to 80.365 (Figure S3A; extended model: 1.35 to 90.93, Figure S4A). As anticipated, the NRMSE values were strongly correlated between the 1^st^ and the 2^nd^ wave experiment (Spearman’s ρ = 0.87, Figure S3B; extended model: Spearman’s ρ = 0.79, Figure S4B), consistent with similar evolutionary dynamics driving the P-element invasion in both EE experiments.

In the top-ranked scenario (Table 2), approximately 20% of the isofemale lines were predicted to carry the P-element (*p*_carrier_ = 0.204; (0.155 – 0.257)), which is slightly below the empirical estimate of 25 – 44% [33]. Additionally, the piRNA cluster size (fraction of each chromosome with P-element-regulatory properties, *f*_regulatory_) under this scenario is 1.7% (1.3% – 2.8%), which is approximately half the size as suggested previously for *D. melanogaster* [37]. We observed similar values for *p*_carrier_ and *f*_regulatory_ in the top-ranked parameter combination for the extended model: *p*_carrier_ = 21.5% (15.3% – 26%) and *f*_regulatory_ = 1.4 % (1.0% – 2.2%) (Table S2).

**Table 2.**
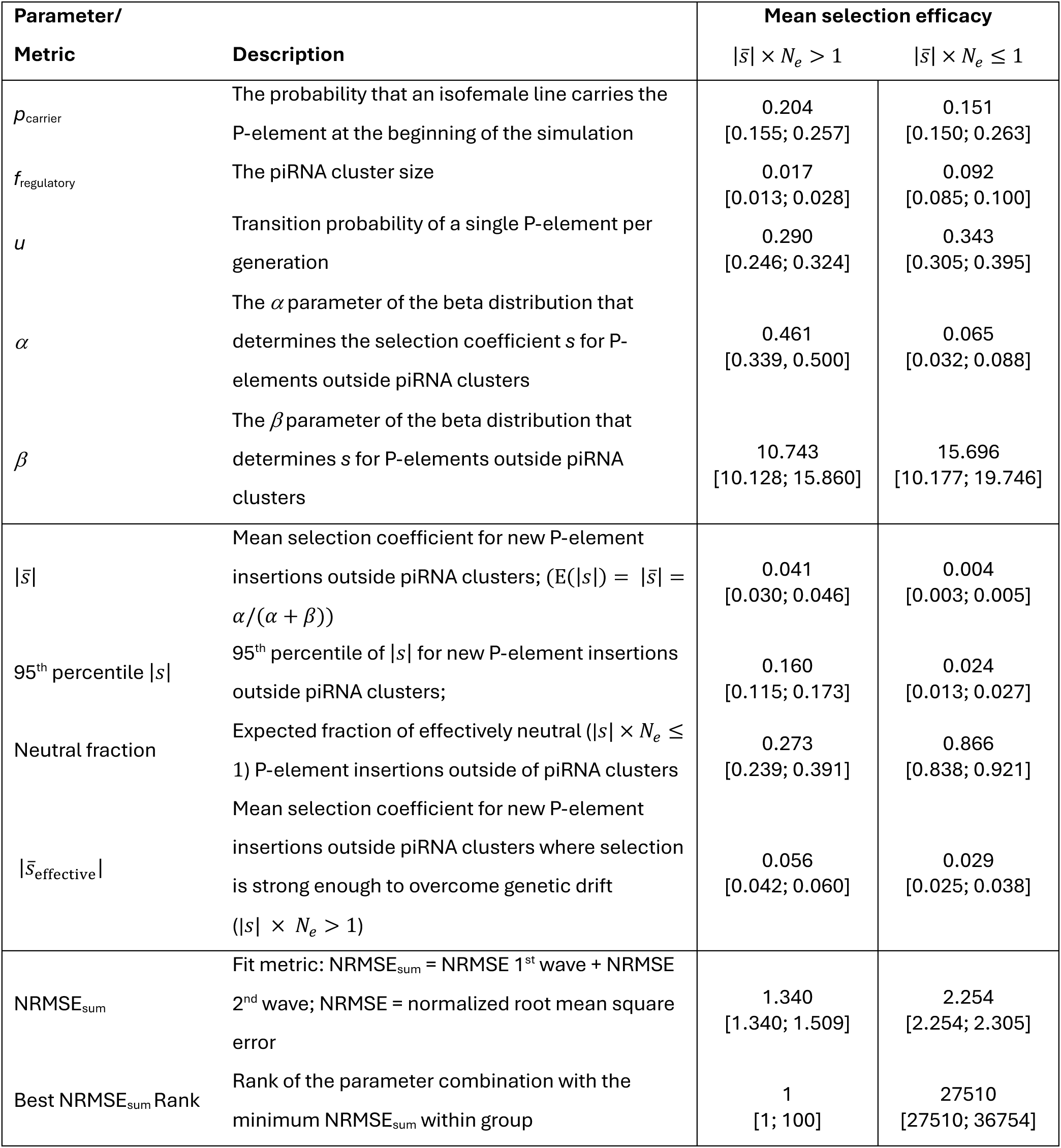
Summary of top-ranked parameter combinations from the parameter exploration with the Gaussian Process model. Parameter combinations are grouped *post hoc* by mean selection efficacy (|*s̅*| × *N*_*e*_) under the assumption of *N*_e_ = 221 [54]. Brackets indicate the range across the top 100 parameter combinations ranked by lowest NRMSE_sum_. See Supplementary Table S2 for a summary from the parameter exploration with the Gaussian Process models trained on simulation outcomes of the extended individual-based simulation model. Results are robust to changes in the effective population size estimate *N*_e_ (Tables S3-4).

| Parameter/<br>Metric |  | Mean selection efficacy |  |
| --- | --- | --- | --- |
| | Description | $\vert \bar{s}\vert \times N_e > 1$ | $\vert \bar{s}\vert \times N_e \leq 1$ |
| $p_{\text{carrier}}$ | The probability that an isofemale line carries the P-element at the beginning of the simulation | 0.204<br>[0.155; 0.257] | 0.151<br>[0.150; 0.263] |
| $f_{\text{regulatory}}$ | The piRNA cluster size | 0.017<br>[0.013; 0.028] | 0.092<br>[0.085; 0.100] |
| $u$ | Transition probability of a single P-element per generation | 0.290<br>[0.246; 0.324] | 0.343<br>[0.305; 0.395] |
| $\alpha$ | The $\alpha$ parameter of the beta distribution that determines the selection coefficient $s$ for P-elements outside piRNA clusters | 0.461<br>[0.339; 0.500] | 0.065<br>[0.032; 0.088] |
| $\beta$ | The $\beta$ parameter of the beta distribution that determines $s$ for P-elements outside piRNA clusters | 10.743<br>[10.128; 15.860] | 15.696<br>[10.177; 19.746] |
| $\vert \bar{s}\vert $ | Mean selection coefficient for new P-element insertions outside piRNA clusters; ($E(\vert s\vert ) = \vert \bar{s}\vert = \alpha/(\alpha + \beta)$) | 0.041<br>[0.030; 0.046] | 0.004<br>[0.003; 0.005] |
| 95<sup>th</sup> percentile $\vert s\vert $ | 95<sup>th</sup> percentile of $\vert s\vert $ for new P-element insertions outside piRNA clusters; | 0.160<br>[0.115; 0.173] | 0.024<br>[0.013; 0.027] |
| Neutral fraction | Expected fraction of effectively neutral ($\vert s\vert \times N_e \leq 1$) P-element insertions outside of piRNA clusters | 0.273<br>[0.239; 0.391] | 0.866<br>[0.838; 0.921] |
| $\vert \bar{s}_{\text{effective}}\vert $ | Mean selection coefficient for new P-element insertions outside piRNA clusters where selection is strong enough to overcome genetic drift ($\vert s\vert \times N_e > 1$) | 0.056<br>[0.042; 0.060] | 0.029<br>[0.025; 0.038] |
| $\text{NRMSE}_{\text{sum}}$ | Fit metric: $\text{NRMSE}_{\text{sum}} = \text{NRMSE 1}^{\text{st}} \text{ wave} + \text{NRMSE 2}^{\text{nd}} \text{ wave}$; NRMSE = normalized root mean square error | 1.340<br>[1.340; 1.509] | 2.254<br>[2.254; 2.305] |
| Best $\text{NRMSE}_{\text{sum}}$ Rank | Rank of the parameter combination with the minimum $\text{NRMSE}_{\text{sum}}$ within group | 1<br>[1; 100] | 27510<br>[27510; 36754] |

For the top-ranked parameter combination, the transposition rate (*u*) was 0.290 (0.246 – 0.324) per P-element per generation (extended model: 0.339 (0.268 – 0.391)). In the extended model, the probability of a clean excision event at the original insertion site (*v*) associated with the top-ranked parameter combination was 0.188 (0.004 – 0.244) per transposing P-element. Empirical estimates of transposition rates *u* for P-elements range between approximately 10^−3^ to 10^−1^ per element per generation [30,51–53], although these values can vary widely depending on genetic background, the presence of repression mechanisms, and environmental factors. We would like to point out that the realized transposition rate in our individual-based simulations is typically lower than the specified *u*, due to purifying selection and the fact that flies with successful P-element repression — triggered via P-element insertions into piRNA clusters — effectively have a transposition rate of zero. Consequently, realized excision probabilities are also reduced in the extended model, as excision events in our model are conditional on transposition events. Given this structure, we reason that the exact value of the excision probability *v* is of secondary importance for the overall invasion dynamics over the observed timescale — a conclusion supported by the wide range of *v* values observed among the 100 top-ranked parameter combinations ((0.044 – 0.244), Table S2).

The parameter combination resulting in the best agreement between predicted and experimental data indicated that a significant amount of purifying selection against new P-element insertions is necessary to account for the observed invasion dynamics. The top-ranked parameter combination — indicated by the lowest NRMSE_sum_ (Table 2) — was obtained for *⍺* = 0.461 and *β* = 10.743, which yielded to an average genome-wide selection coefficient *s* of −0.041(−0.046 – −0.030) for P-element insertions outside piRNA clusters. For the extended model, the top-ranked parameter combination included *⍺* = 0.323 and *β* = 11.766, resulting in an average genome-wide selection coefficient of – 0.027 (−0.044 – −0.018) for new P-element insertions outside piRNA clusters (Table S2). Notably, the top-ranked dominance coefficient *h* was 0.749, yielding an average selection coefficient of −0.02 in heterozygotes — identical to the average heterozygous selection coefficient associated with the top-ranked parameter combination in our original analysis assuming co-dominance (*h* × *s̅* = −0.02). Because fitness effects in the extended model depend on the product of *h* and *s*, metrics based solely on *s* (or *h*) should be interpreted with caution and are not directly comparable to results from the original model assuming co-dominance. Moreover, the values of the dominance coefficient (*h*) from our extended parameter exploration should be interpreted with caution. In the 2^nd^ wave experiment, we assumed complete homozygosity of the individual founder lines: any residual heterozygosity for P-element insertions present in the founding lines would most likely affect the *h* parameter values resulting in the best agreement. Moreover, the next sequenced time point in our experiment was generation 10, when homozygosity would have decreased substantially due to continuous outbreeding and the accumulation of new P-element insertions. Consequently, our experimental data provides limited resolution to distinguish between the effects of *s* and *h* × *s*. Therefore, we assumed co-dominance (*h = 0.5*) in our main analysis.

Overall, our analyses indicate that selection is effective against P-element insertions in experimental *Drosophila* populations, which have typically effective population sizes *N*_e_< 300 [41,54]. Based on the average *N*_e_ estimate of 221 from the 1^st^ wave experiment [54] and the aforementioned top-ranked parameter combination, our results indicate that only 27% (23.9% – 39.1%) of new P-element insertions are effectively neutral (i.e., 2 × *h* × |*s̅*| × *N*_e_ ≤ 1, Table 2; extended model: 37.4 % (23.3% – 40.7%), Table S2) in our experimental populations. Additionally, our findings suggest that strong selection can occur, with the 95^th^ percentile of the selection coefficient value (|*s*|) reaching 0.16 (0.115 – 0.173) assuming co-dominance (Table 2).

We also used the mean *N*_e_ estimate from the 1^st^ wave experiment [54] to categorize parameter combinations into two distinct scenarios: those exhibiting efficient purifying selection against the average P-element insertion, and effectively neural ones. In the latter case, genetic drift is so strong that the invasion trajectory of an average P-element insertion is indistinguishable from a neutral one, even if the insertion is slightly deleterious. Simulation scenarios classified as effectively neutral had the best agreement with the experimental data when they included larger piRNA clusters and higher P-element activity (Table 2, Table S2).

Nevertheless, simulation scenarios that included efficient selection against the majority of P-element insertions showed a better agreement with the empirical invasion dynamics; both in our original analysis assuming co-dominance and in the extended model with variable dominance and excision probability (Figure 6, extended model: Figure S5). Scenarios with mean selection efficacy accurately reproduced both the rapid increase in P-element copy number that occurred during the 1^st^ wave and the transient plateau that was observed between generations 0 and 10 in the 2^nd^ wave. In contrast, scenarios in which the average P-element insertion is effectively neutral consistently failed to replicate these patterns (Figure 6, extended model: Figure S5). This conclusion holds even when potential biases were considered, such as focusing solely on the parameter combination with the lowest NRMSE_sum_. The qualitative pattern remains consistent across the 100 combinations with the lowest NRMSE sums (Figure S6; extended model: Figure S7). Moreover, our results remain consistent when the *N*_e_ estimate was modified, as this parameter serves as a threshold for mean selection efficacy in our approach. Doubling the *N*_e_ estimate from 221 to 442 did not result in any qualitative alterations to the outcomes, thereby reinforcing the conclusion that purifying selection plays a pivotal role in shaping the observed P-element invasion dynamics (Table S3, extended model: Table S4).

**Figure 6.**
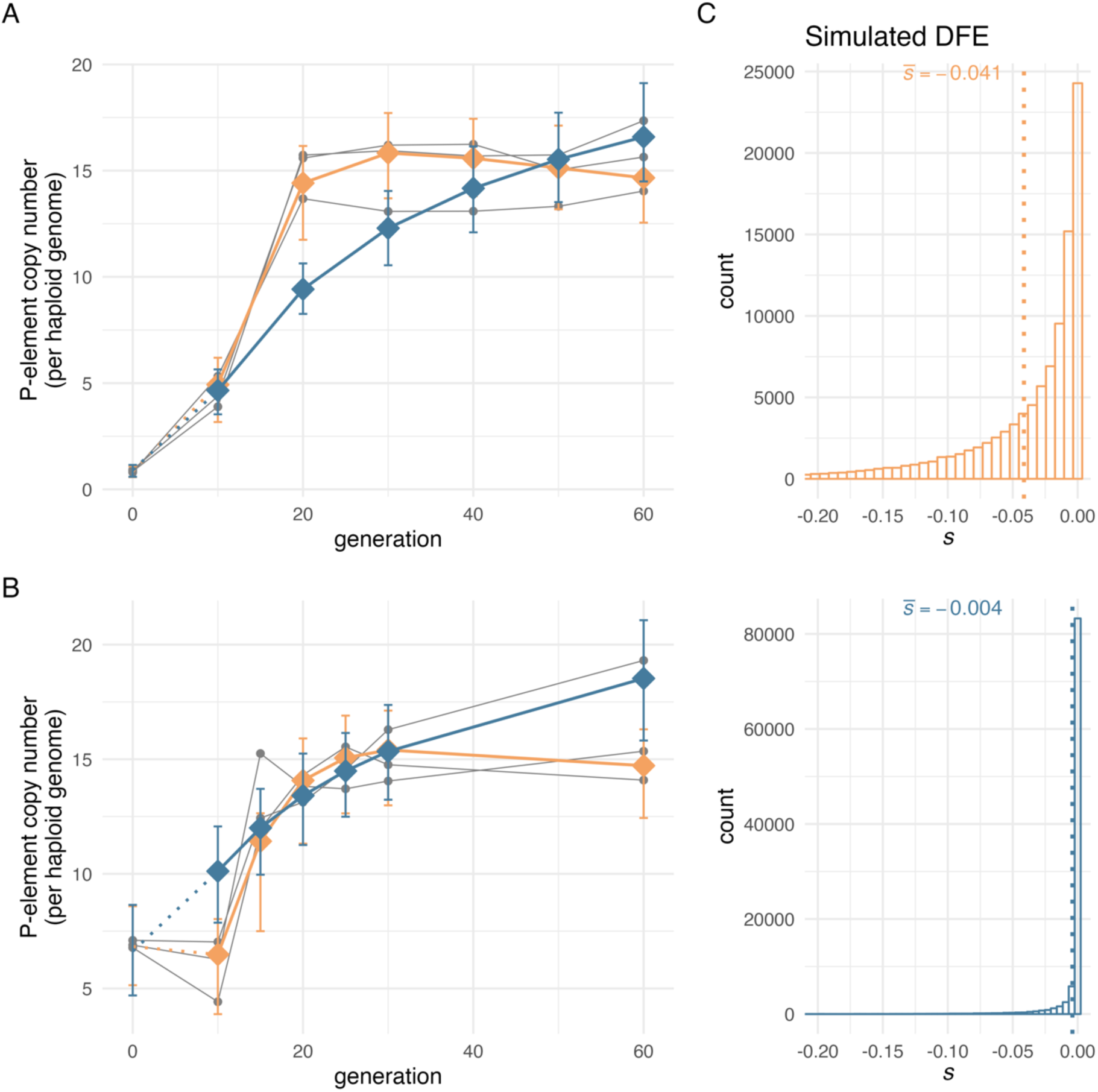
Gaussian Process (GP) predictions provide a good fit to the empirical data — the top-ranked parameters were identified using 1^st^ and 2^nd^ wave data. GP predictions from the parameter combinations with the lowest NRMSE sum when compared against empirical data are shown for scenarios without mean selection efficacy (|*s̅*| × *N*_*e*_ ≤ 1, steel blue) and scenarios with mean selection efficacy (|*s̅*| × *N*_*e*_ > 1, amber), *N_e_* = 221 [54]. Each grey line represents an empirical evolution replicate, with sequenced time points indicated by dots. GP predictions are indicated by connected colored diamonds. The error bars show the range between the 2.5^th^ and 97.5^th^ percentiles of P-element copy number trajectories simulated with the individual-based model. These simulations were run with the same parameters as those used for GP prediction (Table 2). The error bars are shown at the same time points where empirical data were sequenced. Note that GP predictions do not cover generation 0; instead, predictions from generation 10 are connected to the individual-based simulation averages at generation 0 to aid visual interpretation. (A) 1^st^ wave experiment. (B) 2^nd^ wave experiment. (C) Simulated distribution of fitness effects (DFE) used in (A) and (B). Due to selection, the actual distribution of selection coefficients for segregating P-element insertions in the simulated populations will be skewed toward 0. Results are robust across the 100 parameter combinations with the lowest NRMSE sums (Figure S6). See Supplementary Figure S5 for predictions from the GPs that were trained on simulation outcomes of the extended individual-based model.

## Discussion

### The power of Experimental Evolution to study TE dynamics

To evaluate the role of purifying selection in controlling TEs, we investigated the dynamics of P-element invasions in two *Drosophila* EE experiments. Because low-frequency TE insertions, a hallmark of the early stages of an invasion process, are very difficult to study empirically, we used the P-element copy number as a summary statistic to describe the invasion dynamics at the population level. Through careful model analysis, we estimated the distribution of fitness effects of P-element insertions in these experiments.

While beneficial fitness effects have been documented for some TE insertions (e.g., [55–59]), we reasoned that such cases are rare, and thus unlikely to substantially impact the population-level dynamics of a P-element invasion. Furthermore, a previous analysis of the 1^st^ wave experiment and further replicate populations that were set up simultaneously from the same founder population in May 2011 found no evidence for positive selection acting on any P-element insertion site over 60 generations [41]. We thus focused our analysis on deleterious effects of P-element insertions. We calibrated an individual-based simulation model to data from the two EE studies to estimate the distribution of deleterious fitness effects, first from the 1^st^ wave alone, and then from both the 1^st^ and 2^nd^ waves combined. Both approaches yielded similar results. Remarkably, top-ranked parameter combinations identified only from the 1^st^ wave data accurately predicted the invasion dynamics observed in the 2^nd^ wave experiment, conducted four years later under different initial conditions. Overall, we observed that substantial purifying selection outside of piRNA clusters (*s̅* = −0.041 (−0.046 – −0.030)) was necessary to accurately capture the observed P-element invasion patterns in both experiments. TE invasions are challenging to analyze in natural populations due to their rapid dynamics and confounding demographic factors such as gene flow. Excitingly, this study not only demonstrates that purifying selection is a key evolutionary force shaping P-element invasions, but also highlights the utility of EE, in combination with simulation and modeling-based approaches, for studying TE invasions in the lab.

It is important to note, however, that our results are constrained by the *a priori* chosen parameter ranges and combinations in our analyses. It remains possible that unexplored regions of the parameter space could yield even better agreement between GP predictions and experimental data. However, grouping parameter combinations *post hoc* by their mean selection efficacy clearly demonstrates that efficient purifying selection against the majority of P-element insertions is consistently associated with better agreement between simulations and experimental data. Moreover, our analysis of the extended model — which allows for variation in dominance coefficients and excision probabilities — leads to similar conclusions based on an independently generated parameter set. Thus, although our parameter exploration approach is heuristic in nature, it robustly demonstrates that purifying selection plays a key role in shaping P-element invasion dynamics in experimental *D. simulans* populations.

In our analysis, we grouped parameter combinations *post hoc* based on the mean selection efficacy (2 × *h* × |*s̅* | × *N*_*e*_). We recognize that this approach is a simplification: in reality, evolutionary dynamics are shaped by genome-wide selection, and selection can act effectively across many loci even when individual P-element insertions have negligible fitness effects. Nevertheless, using the mean selection efficacy provides a useful heuristic to capture overall trends across parameter combinations, and allows us to visually demonstrate that weak selection against the majority of P-element insertions is insufficient to explain the observed invasion dynamics in our experimental populations.

### Alternative mechanisms of TE regulation

A key feature of our study is the individual-based modeling framework used to simulate P-element invasion dynamics in experimental *Drosophila* populations. As with any model, it is based on a number of simplifying assumptions that inevitably omit some aspects of biological realism, and these assumptions might influence the model outcomes. For example, our individual-based simulations are based on the classic trap model, which posits that a single TE insertion in a piRNA cluster silences all TEs of that type. Given that the distribution of TEs in piRNA clusters differs from predictions under the trap model [60,61] and that the removal of piRNA clusters does not necessarily induce TE activity [62], alternative mechanisms of TE silencing must be considered.

One potential mechanism could be paramutations, whereby insertions outside of piRNA clusters can be converted into piRNA-producing loci in the presence of maternally deposited piRNAs [63–65]. In a recent study, Scarpa *et al*. (2023) demonstrated with comprehensive computer simulations that this additional layer of defense against TEs can help to resolve some of the discrepancies between observed and predicted TE copy numbers under the classic trap model in natural *Drosophila* populations [63]. Since Scarpa *et al.* (2023) focused on the overall abundance of TE families in *D. melanogaster*, it remains unclear to what extent paramutations are shaping TE invasion processes in natural and experimental populations over time. Given the excellent fit between the simulations under the trap model and our empirical data, it is possible that paramutations are not essential to explain the observed P-element dynamics in our two EE experiments. Even more importantly, the transient P-element copy number plateau observed between generations 0 and 10 in the 2^nd^ wave experiment, along with the subsequent increase in copy number in later generations, is more readily explained by purifying selection rather than by paramutations. Further work is required to evaluate the extent to which paramutations influence the invasion dynamics of the P-element and other TEs.

### Most TE insertions are deleterious

The effective population size is a key factor in determining whether a mutation is subject to selection or whether its fate is primarily driven by genetic drift [66]. In EE experiments with *Drosophila*, such as those presented in this work, estimates of effective population size typically range from 200 to 400 [41,54]. At this effective population size, purifying selection can only act on mutations with strongly deleterious effects. In natural *Drosophila* populations, the effective population size is several orders of magnitude larger [67,68]. This implies that in natural *Drosophila* populations, only a small fraction of new P-element insertions is likely to be effectively neutral — much fewer than the 27% (23.9% – 39.1%) estimated from our experimental data. This idea aligns well with the observation that only a small fraction of the genome shares P-element insertions between *D. melanogaster* and *D. simulans* [35]. While this is typically attributed to non-random insertions of actively transposing P-elements [12], we propose that purifying selection in large natural populations might drive independent enrichment of P-element insertions at genomic regions subject to relaxed purifying selection.

## Conclusions

Our study highlights the crucial role of purifying selection in shaping TE invasion dynamics, specifically in the context of P-element invasions in *D. simulans*. Using an individual-based model, we demonstrated that substantial purifying selection is needed to replicate the observed invasion dynamics in two EE studies, assuming that each P-element insertion is either neutral or detrimental, but not beneficial [41]. This study not only highlights EE as a valuable tool for understanding TE behavior, but also illustrates how computational modeling can complement experimental work to study TE invasions. Future work could expand this modeling framework to include additional regulatory mechanisms such as paramutations, advancing our understanding of TE behavior in diverse environments.

## Methods

### Experimental Evolution

The details of the 1^st^ wave EE experiment have been previously published [22,54]. While ten evolutionary replicates were set up simultaneously from the same founder population in May 2011, we focus on three replicate populations for which detailed P-element analyses have already been conducted and published [54], and we directly use the P-element copy number estimates reported in Kofler *et al.* (2022). Additionally, selecting three replicates ensures a balanced number of replicates between the 1^st^ and 2^nd^ wave experiments, which is important when calculating the agreement between GP predictions and experimental data (please refer to the “Parameter exploration with Gaussian Process surrogate models” section for further details).

Pool-Seq data for generations 0 and 10 of the 2^nd^ wave EE experiment are available in Langmüller *et al.* (2023) [11]. This study adds generation 15 – 60 of the 2^nd^ wave experiment. This section only provides an overview of the experiments and specific information regarding already published data can be found in those sources. To maintain consistency and facilitate comparison between studies, we used the same replicate identifiers as in the original publications.

#### Setup G maintenance of experimental *D. simulans* populations

For the 1^st^ wave [22], replicate populations were established from 202 isofemale lines collected from a natural *D. simulans* population in Tallahassee, Florida [69], which experienced a P-element invasion at the time point of sampling [35]. For the 2^nd^ wave experiment, three replicate populations were established using the surviving 191 out of 202 isofemale lines that were originally collected [11]. All replicate populations were maintained with non-overlapping generations in a cycling hot environment (12 hours of light at 28°C; 12 hours of darkness at 18°C) and at a constant population size (*N* = 1,000 for the 1^st^ wave; *N* = 1,250 for the 2^nd^ wave) for 60 generations.

#### Genomic sequencing

All experimental populations were sequenced using the Pool-Seq approach [42] with at least 500 flies per sample. In the 1^st^ wave, populations were sequenced every 10^th^ generation, resulting in 21 samples (7 time points; 3 replicate populations) [22,54]. To more effectively monitor the rapid invasion of the P-element, we increased the number of sequenced time points at the beginning of the 2^nd^ wave experiment. For this wave, populations were sequenced at generation 0, as well as at generations 10, 15, 20, 25, 30, and 60, also resulting in 21 samples. Genomic DNA for all new samples was extracted from half the number of individuals that had contributed to the next generation (approximately 600 flies). To facilitate comparison across studies, we adopt the Fxx nomenclature for sequencing samples, where ‘xx’ represents the generation in our EE experiment. Paired-end libraries for generation F60 were prepared and sequenced along with those for generation F0 using the NEBNext Ultra II DNA Library Prep Kit (New England Biolabs, Ipswich, MA) with an insert size of 300bp. Paired-end libraries for generations F15, F20, F25, and F30 were processed in the same way as those for generation F10, using a protocol with 10% of the reagents provided in the NEBNext Ultra II FS DNA Library Prep Kit (New England Biolabs, Ipswich, MA). Steps for all library preparations are described in more detail in [11]. We sequenced all libraries using a 2 × 150 bp protocol on a HiSeq X Ten.

#### P-element copy number as a summary statistic for invasion dynamics

Given the rapid spread of the P-element in our experimental populations, most individual P-element insertions are expected to remain at low frequency. This poses a major challenge for accurately estimating empirical P-element insertion frequencies, irrespective of whether pools of individuals or single individuals are being analyzed. Therefore, we used the estimated P-element copy number per haploid genome as a summary statistic for describing the P-element invasion in the experimental populations. We used DeviaTE [43] to estimate P-element copy number per haploid genome in both EE experiments. DeviaTE is a bioinformatics tool that estimates TE abundance from Illumina sequencing data. It maps sequencing reads to both TE consensus sequences and a user-defined set of single-copy genes. DeviaTE calculates P-element copy number by comparing the read coverage of the P-element consensus sequence with the mean coverage of the single copy genes, while accounting for the presence of internal and terminal deletions within individual P-element copies [43,70]. For the 1^st^ wave, P-element copy number estimates were calculated by Kofler *et al.* 2022 using the mean coverage of three single copy genes (*rpl32, tra=ic jam, rhino*) as normalization factor. These estimates were previously published in Kofler *et al.* 2022 and are available in the supplemental material (replicates 1, 3, and 5 from the hot environment) [54].

For 2^nd^ the wave, we used ReadTools (v1.5.2.) [71] to demultiplex *(–maximumMismatches 3 –maximumN 2*) and trim barcoded sequencing files based on sequencing quality (*– mottQualityThreshold 15 –disable5pTrim –minReadLength 25*). Since DeviaTE does not take paired-end read information into account [43], we merged paired-end read files prior mapping. We mapped reads to a reference genome that included the *Drosophila* P-element consensus sequence [72] along with three single copy genes (*rpl32, tra=ic jam, rhino),* using bwa bwasw (v0.7.17; *–M*) [73]. Finally, we estimated P-element copy number per haploid genome with DeviaTE (*–minID 1*) [43].

Our genomic analysis pipeline follows Kofler *et al*. 2022 with some minor differences (e.g., we used a more recent version of bwasw) [54]. To verify the consistency between pipelines, we re-analyzed three randomly selected samples from Kofler *et al*. (2022) using our updated pipeline. The copy number estimates from the two pipelines differed by less than 5 %, which we considered an acceptable level of consistency (data not shown).

### P-element invasion models

#### Individual-based models

We implemented an individual-based P-element invasion model in SLiM (v4.0.1) [46]. Our individual-based model simulates an obligatory outcrossing population with discrete, non-overlapping generations and a constant census size of 1,000 individuals (flies). Figure 3 provides a schematic overview of the individual-based simulation model.

Each fly has a diploid genome consisting of five chromosomes, each 32.4 Mb in length, resulting in a total genome size of 162 Mb, as estimated for *D. simulans* by flow cytometry [74]. We modeled a constant recombination rate of 4×10^−8^ per bp per generation, based on the *Drosophila simulans* recombination map [75]. Unlike its sister species *D. melanogaster*, *D. simulans* lacks large segregating chromosomal inversions [69,76]. Additionally, only a small proportion of the genome near centromeric regions experiences suppressed recombination in *D. simulans* [77]. Given these characteristics, assuming a uniform recombination rate across the genome is a reasonable approximation in our individual-based model. Moreover, previous simulation studies have shown that recombination rate variation has only a minor effect on simulated TE invasion dynamics mimicking similar experimental settings [44].

At the end of each chromosome, we modeled a piRNA cluster region capable of regulating P-element activity [37,44]. The size of simulated piRNA clusters is regulated by the model parameter *f*_regulatory_. The only genomic variants included in the individual-based model are P-element insertion sites (Figure 3A).

The ancestral population (generation 0) is initialized by mixing 200 inbred fly strains (isofemale lines), using five flies per line. Single lines might be P-element carriers, with a user-specified probability *p*_carrier_ (Table 1). If a line carries the P-element, the number of initial P-element insertion sites is drawn from a Poisson distribution, where the rate parameter λ is chosen such that, given the value of *p*_carrier_, the average P-element copy number per haploid genome for the simulated EE experiment fits the empirical data (1^st^ wave: 0.86 P-element copies per haploid genome; 2^nd^ wave: 6.92). P-element insertion sites are chosen randomly across the genome and all flies from the same isofemale line are isogenic, meaning they share the same initial set of P-element insertion sites. For the 1^st^ wave, we modeled heterozygous initial P-element insertions, because the P-element invasion was recent [35]. For the 2^nd^ wave, we modeled homozygous initial P-element insertions, given that the isofemale lines had been maintained at small population sizes for 4.5 years [11], causing increased inbreeding and a likely establishment of defense mechanisms against further P-element proliferation.

We acknowledge that the assumptions of isogenic flies and complete homozygosity for the 2^nd^ wave likely oversimplify the genetic structure of isofemale lines. However, given the lack of detailed, time-resolved data on the precise inbreeding state of the isofemale lines, which have undergone extreme bottlenecks and have experienced overlapping generations, we chose to model complete homozygosity for the founder lines of the 2^nd^ wave.

To account for a host defense mechanism against the P-element, we incorporated the trap model into our individual-based model [39,40]. Under the trap model, P-element activity is regulated by insertions into piRNA clusters [37]. In our individual-based model, single P-elements transpose with probability *u* per generation, unless a fly carries at least one P-element insertion in a piRNA cluster. Note that this rule applies also for the isofemale lines: any isofemale line that harbors a piRNA-cluster insertion at generation 0 also has zero transposition activity. We chose the range of *u* (0.15 – 0.5, Table 1) based on the observed invasion dynamics in the 1^st^ wave experiment. While other experimental studies suggest that individual P-element copies transpose at rates ranging from 10^−3^ to 10^−1^ per element per generation [30,51–53], these effective transposition rates (*u’*) depend on additional factors such as selection, and the frequency of active elements.

We estimated *u’* from the first 10 generations of the 1^st^ wave experiment. Assuming a copy-and-paste mechanism (i.e., excision probability = 0) and minimal host suppression at the beginning of the P-element invasion, we expect that the P-element copy number grows exponentially with time:

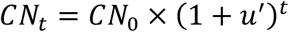

where *CN_t_* is the P-element CN at time t, *CN_0_* is the initial copy number, and *u’* is the effective transposition rate per generation. For the early stages of the 1^st^ wave experiment, we estimated effective transposition rates *u’* between 0.16 and 0.21, depending on the replicate. This suggests that, in the absence of selection *x* against individual P-element insertions, the transposition probability *u* must be at least 0.16, as *u’ = u – x.* The upper bound of *u = 0.5* was chosen to ensure that effective transposition rates remain positive (*u’ > 0*) even under strong purifying selection (x > 0).

In our main analysis, we assumed that P-element proliferation operates exclusively via a copy-and-paste mechanism, with no excision events. This simplification is based on evidence from previous studies that P-elements can increase their copy number through mechanisms such as sister chromatid-mediated gap repair following excision [78] and transposition into unreplicated DNA during cell division, which facilitates copy number increase without significant loss of existing elements [12].

However, in our extended model (Figure S1), each transposing P-element has a probability *v* of being excised from its original insertion site. If a P-element insertion is homozygous, transposition and excision are handled independently for each P-element copy. If a single P-element insertion occurs in any piRNA cluster, all P-elements within the whole genome are rendered inactive immediately and are unable to transpose further (Figure 3B).

We assumed that P-element insertions within piRNA clusters are neutral (selection coefficient *s* = 0). For all other P-element insertions, the absolute value of *s* is drawn from a beta distribution with shape parameters α and β (Figure 3C). We considered only purifying selection against P-elements (*s* <= 0), because no signature of adaptive selection was detected in the 1^st^ wave experiment and further replicate populations that were set up simultaneously from the same founder population in May 2011 [41]. In our main analysis, we assumed co-dominant fitness effects by setting the dominance coefficient (*h*) to 0.5. In the extended model, *h* is treated as an additional model parameter that can vary between simulation runs, with values ranging between 0 and 1 (Table S1). Individuals with no P-element insertions at a single locus have a fitness of 1, heterozygous individuals have a fitness of 1 +*hs*, and individuals homozygous for the P-element insertion have a fitness of 1 + *s*, where *s* is always <= 0. With more than one P-element insertion site, the fitness effects from each site are combined with the others multiplicatively.

In the basic model (main text), we systematically varied five parameters: the probability that a single line carries the P-element, the fraction of the chromosome capable of triggering a defense against the P-element (piRNA cluster size), the transposition probability, and two parameters governing the distribution of selection coefficients (Table 1). For the extended model, we additionally varied the excision probability *v* and the dominance coefficient *h*, resulting in a total of seven parameters (Table S1).

For both individual-based simulation models, we repeated simulations 100 times for each parameter combination, and recorded the average P-element copy number per haploid genome at the same time points used in the empirical EE experiments (Figure 3D). This approach allowed us to directly compare simulation outcomes with observed P-element copy numbers.

#### Gaussian Process surrogate models

Because our individual-based model has at least five parameters that can be varied (Table 1; extended model: seven parameters, Table S1), exploring the full range of this parameter space with the individual-based model would be a complex and time-intensive task [47]. To streamline this process, we employed statistical emulation, replacing the individual-based model with a surrogate model that can rapidly and accurately predict the individual-based model’s behavior [79]. GP surrogate models [48,80] are particularly well-suited for this task because they are non-parametric models that define distributions over functions. This characteristic allows them to efficiently extrapolate between sparsely sampled data points. For a detailed introduction to GPs, we defer to Rasmussen and Williams [48].

We implemented our GPs in Python (v3.10.6) using the GpyTorch (v1.11) [81] and PyTorch (v2.0.1) [82] libraries for efficient GP modeling. We trained a separate GP for each EE experiment (1^st^ and 2^nd^ wave) and for each individual-based model type (basic model with five parameters; extended model with seven parameters), resulting in a total of four GPs. Specifically, we used a multi-task GP [81,83] to model the relationship between six tasks — the P-element copy number for the evolved generations of the corresponding EE experiment — and five model parameters (Table 1; extended model: seven parameters, Table S1), which serve as predictors across all tasks. Due to time-related dependencies of P-element copy numbers within one experiment, we assumed that tasks are correlated and modeled their relationship using a rank-1 covariance structure. This means that any correlation between single tasks is determined by one single latent factor. The multi-task GP consists of a constant mean function providing a baseline P-element copy number estimate for each task (constant mean = 0 copies), and a covariance function that accounts for both: input-dependent correlation within one task (*k*_input_) and correlations across tasks (*k*_task_):

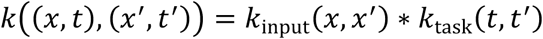

Where *k*_input_(*x*, *x*′) is a radial basis function kernel modeling the covariance between two input data points *x* and *x’* [48], and *k*_task_(*t*, *t*′) models the relationship between tasks *t* and *t’* [81,83]. Using a rank-1 task-kernel simplifies the task covariance matrix *K*_task_ to:

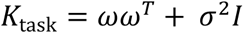

Where ω is a vector of task weights and *σ*^2^ represents task-specific, independent noise. During GP training, the GP “learns” the task weights ω, *k*_input_ hyper-parameters, and task-specific noise by maximizing the log marginal likelihood over observed training data.

We trained GPs on a dataset of 1000 points (extended model: 2500 points) using Latin Hypercube Sampling (LHS), a method that ensures evenly distributed exploration of the entire input domain (basic model: Table 1; extended model: Table S1), where P-element copy number time series were generated with the experiment-specific individual-based model. For the basic model, we used the Adam optimizer from PyTorch with a learning rate of 0.01 [82] to train each GP over 40 rounds (50 iterations per round). For the extended model, which includes two additional parameters and therefore required more training, we trained each GP for 50 rounds, with 100 optimization iterations per round using the same optimizer and learning rate. After each training round, we saved a snapshot of the GP and evaluated it against an independent validation dataset consisting of 5000 LHS points (extended model: 10000 LHS points) to determine the optimal training duration and avoid overfitting. We selected the GP snapshot with the lowest root mean square error (RMSE) on the validation dataset for further analysis (basic model: snapshot 18 for 1^st^ wave, snapshot 19 for 2^nd^ wave; extended model: snapshot 36 for 1^st^ wave, snapshot 44 for 2^nd^ wave).

To evaluate the GP’s performance, we predicted P-element copy number time series for a test dataset consisting of 5000 LHS data points (extended model: 10000 LHS points). To make GP model performance comparable between the two EE experiments, we report RMSE values normalized by the average observed P-element copy number per time point. This normalization was necessary because the 2^nd^ wave, which has higher absolute P-element copy numbers in the ancestral generation, would otherwise naturally result in higher RMSEs. The trained GP was only challenged when predicting large P-element copy numbers in the test data.

This issue is likely due to our choice of a constant mean of 0 across all tasks (P-element copy number per generation). When data points are sparse, the GP reverts to the prior mean of 0, which tends to downwardly bias the predictions (an alternative approach would be to set the mean to the average observed value in the training data for each respective generation). Another potential source of error lies in the kernel choice. Simulated P-element invasions often involve rapid changes, and the RBF kernel we used might not always fully capture these rapid dynamics. Alternative kernels, such as the Matérn kernel [48,84], might be more effective at modeling abrupt shifts in behavior, offering a better fit for this type of system. Additionally, the GP’s Bayesian nature offers a way to improve its predictive power. In Bayesian frameworks, predictions are not single values but are accompanied by uncertainty estimates. This uncertainty can guide further sampling in regions where the GP’s predictions are less confident, a technique known as active learning [84]. Because the trained GPs predicted P-element copy numbers within the empirical range with high accuracy, we did not explore alternative mean function, kernel choices, and active learning strategies in this study.

#### Parameter exploration with Gaussian Process surrogate models

We used our trained GPs to comprehensively explore simulated P-element dynamics across the whole input domain (Table 1; extended model: Table S1). For each EE experiment (1^st^ and 2^nd^ wave), we predicted P-element copy number time series for 10^6^ LHS data points (extended model: 10^7^ LHS points). The GPs allowed for rapid calculations: predicting P-element invasion curves for 10^6^ data points took only 2 seconds on a local machine with an i5-12600K CPU and a GeForce RTX4090 GPU. In comparison, running 10^8^ individual-based simulation runs for the same type of analysis would take several days on a computing cluster, depending on available resources.

To identify the parameter values that best explain the observed P-element invasions, we assessed the fit between the predicted and observed P-element copy number time series. For each evolved generation, we computed the RMSE between the GP prediction and the three replicate experimental populations. Next, we normalized the RMSE by dividing by the average observed P-element copy number at the corresponding generation. Finally, we summed the normalized RMSE across all six evolved generations to generate a single error metric between the predicted and empirical P-element invasion curves for each EE experiment. The fit metric across both EE experiments — NRMSE_sum_ — is simply the sum of the NRMSEs for the 1^st^ and 2^nd^ wave experiments. A lower NRMSE value indicates a better agreement between the predicted and observed P-element invasion dynamics.

## Declarations

### Ethics approval and consent to participate

Not applicable

### Consent for publication

Not applicable

### Availability of data and materials

Information regarding data availability and processing for the 1^st^ wave can be found in Kofler *et al.* 2018 and Kofler *et al.* 2022. For the 2^nd^ wave, raw reads of generations 0 and 10 have been previously published (Langmüller *et al*. 2023) and are available from the European Nucleotide Archive (ENA) under project accession number PRJEB54573 (samples SAMEA110271575 – SAMEA110271580). Raw reads of the remaining generations of the 2^nd^ wave experiment are available from ENA under project accession number PRJEB83416. Time-resolved P-element copy number estimates in the experimental populations, our individual-based P-element invasion models implemented in SLiM [46], simulated data, trained GPs, as well as a Jupyter notebook demonstrating the usage of the GPs are available on GitHub at https://github.com/AnnaMariaL/EE-P_element.

### Author’s contributions

CS designed and supervised the experimental evolution study. AML was responsible for the modeling aspects of the study. VN was responsible for the molecular work. BCH contributed to the design of the individual-based simulation model. AML and CS wrote the initial manuscript draft. All authors revised the manuscript and approved the final version.

## Acknowledgements

We thank all current and former members of the Institute of Population Genetics, who contributed to the maintenance of the experimental fly populations over many years. Special thanks to Marlies Dolezal for helpful discussions. We also thank all members of the Messer lab for feedback and support. Special thanks to Meera Chotai, Isabel Kim, and Kiran Chandrasekher for technical support; and last but not least to Sam Champer for technical support and for generously running some GP predictions on his personal computing resources.

## Supplementary Information

### Supplementary Tables

**Table S1.** Model parameters of the extended individual-based P-element invasion model and their considered ranges, including additional model parameters for excision probability (*v*) and dominance (*h*). The range for *p*_carrier_ was guided by a previous study estimating that 25 – 44% of the isofemale lines used in the 1^st^ and 2^nd^ wave experiment carried the P-element [33]. The range for *f*_regulatory_ was based on the estimate that 3.5% of the *Drosophila melanogaster* genome consists of piRNA clusters with TE-regulatory properties [37]. However, because piRNA clusters are challenging to assemble and compare across species [49] and because a previous simulation study suggests that as little as 0.2% may be sufficient for TE control [50], we allowed *f*_regulatory_ to vary rather than fixing it. The range for *u* was informed by the observed P-element invasion dynamics in the 1^st^ wave experiment. Specifically, the lower bound of 0.15 was chosen based on effective transposition rate estimates derived from generations 0 and 10 in the 1^st^ wave experiment. The upper bound of 0.5 ensures that effective transposition remains possible even under strong purifying selection scenarios.

| Parameter | Description | Range |
| --- | --- | --- |
| $p_{\text{carrier}}$ | The probability that an isofemale line carries the P-element at the beginning of the simulation | [0.15, 0.5] |
| $f_{\text{regulatory}}$ | piRNA cluster size: the fraction of the chromosome with P-element-regulatory properties (once at least one P-element insertion is acquired) | [0.01, 0.1] |
| $u$ | The probability of transposition of a single P-element per generation | [0.15, 0.5] |
| $v$ | The probability of a clean excision event at the original insertion site, given that a P-element transposes | [0, 0.25] |
| $\alpha$ | The shape parameter $\alpha$ of the beta distribution that defines the selection coefficient $s$ for P-element insertions outside piRNA clusters | [0.001, 0.5] |
| $\beta$ | The shape parameter $\beta$ of the beta distribution that defines the selection coefficient $s$ for P-element insertions outside piRNA clusters | [10, 20] |
| $h$ | The dominance coefficient specific to each P-element insertion | [0, 1] |

**Table S2.** Summary of top-ranked parameter combinations from the parameter exploration with the Gaussian Process model trained on the extended individual-based P-element invasion model that includes additional parameters for excision probability (*v*) and dominance (*h*). Parameter combinations are grouped *post hoc* by mean selection efficacy (2 × *h* × |*s̅*| × *N*_*e*_) under the assumption of *N_e_* = 221 [54]. Brackets indicate the range across the top 100 parameter combinations ranked by lowest NRMSE_sum_. Variation among the top 100 parameter combinations reflects that agreement between GP predictions and experimental data depends on specific combinations of model parameters, and that the fitness effects of new P-element insertions arise from the product of *h* × *s*, limiting the interpretability of *s* or *h* individually.

| Parameter/<br>Metric | Description | Mean selection efficacy |  |
| --- | --- | --- | --- |
| | | $2 \times h \times \vert \bar{s}\vert \times N_e > 1$ | $2 \times h \times \vert \bar{s}\vert \times N_e \leq 1$ |
| $p_{\text{carrier}}$ | The probability that an isofemale line carries the P-element at the beginning of the simulation | 0.215<br>[0.153; 0.260] | 0.163<br>[0.150; 0.228] |
| $f_{\text{regulatory}}$ | The piRNA cluster size | 0.014<br>[0.010; 0.022] | 0.083<br>[0.074; 0.092] |
| $u$ | Transition probability of a single P-element per generation | 0.339<br>[0.268; 0.391] | 0.347<br>[0.316; 0.428] |
| $v$ | Excision probability at the original insertion site given that a P-element transposes | 0.188<br>[0.004; 0.244] | 0.036<br>[0.000; 0.182] |
| $\alpha$ | The $\alpha$ parameter of the beta distribution that determines the selection coefficient $s$ for P-elements outside piRNA clusters | 0.323<br>[0.284; 0.499] | 0.385<br>[0.107; 0.498] |
| $\beta$ | The $\beta$ parameter of the beta distribution that determines $s$ for P-elements outside piRNA clusters | 11.766<br>[10.072; 19.976] | 10.785<br>[10.135; 19.513] |
| $h$ | Dominance coefficient | 0.749<br>[0.488; 0.967] | 0.066<br>[0.032; 0.362] |
| $\vert \bar{s}\vert $ | Mean selection coefficient for new P-element insertions outside piRNA clusters; ($E(\vert s\vert ) = \vert \bar{s}\vert = \alpha/(\alpha + \beta)$) | 0.027<br>[0.018; 0.044] | 0.034<br>[0.006; 0.042] |
| 95<sup>th</sup> percentile $\vert s\vert $ | 95<sup>th</sup> percentile of $\vert s\vert $ for new P-element insertions outside piRNA clusters; | 0.119<br>[0.079; 0.165] | 0.143<br>[0.035; 0.163] |
| Neutral fraction | Expected fraction of effectively neutral ($2 \times h \times \vert s\vert \times N_e \leq 1$) P-element insertions outside of piRNA clusters | 0.374<br>[0.233; 0.407] | 0.696<br>[0.684; 0.822] |
| $\vert \bar{s}_{\text{effective}}\vert $ | Mean selection coefficient for new P-element insertions outside piRNA clusters where selection is strong enough to overcome genetic drift ($2 \times h \times \vert s\vert \times N_e > 1$) | 0.042<br>[0.029; 0.058] | 0.094<br>[0.031; 0.133] |
| NRMSE<sub>sum</sub> | Fit metric: NRMSE<sub>sum</sub> = NRMSE 1<sup>st</sup> + NRMSE 2<sup>nd</sup> wave<br>NRMSE = normalized root mean square error | 1.353<br>[1.353; 1.418] | 2.196<br>[2.196; 2.266] |
| Best NRMSE<sub>sum</sub> Rank | Rank of the parameter combination with the minimum NRMSE<sub>sum</sub> within group | 1<br>[1; 100] | 123217<br>[123217; 176086] |

**Table S3.** Summary of top-ranked parameter combinations from the parameter exploration with the Gaussian Process model. Parameter combinations are grouped *post hoc* by mean selection efficacy (|*s̅*| × *N*_*e*_) under the assumption of N_e_ = 442 and co-dominance. Brackets indicate the range across the top 100 parameter combinations ranked by lowest NRMSE_sum_.

| Parameter/<br>Metric | Description | Mean selection efficacy |  |
| --- | --- | --- | --- |
| | | $\vert \bar{s}\vert \times N_e > 1$ | $\vert \bar{s}\vert \times N_e \leq 1$ |
| $p_{\text{carrier}}$ | The probability that an isofemale line carries the P-element at the beginning of the simulation | 0.204<br>[0.155; 0.257] | 0.151<br>[0.151; 0.497] |
| $f_{\text{regulatory}}$ | The piRNA cluster size | 0.017<br>[0.013; 0.028] | 0.098<br>[0.092; 0.100] |
| $u$ | Transition probability of a single P-element per generation | 0.290<br>[0.246; 0.324] | 0.343<br>[0.308; 0.367] |
| $\alpha$ | The $\alpha$ parameter of the beta distribution that determines the selection coefficient $s$ for P-elements outside piRNA clusters | 0.461<br>[0.339; 0.500] | 0.039<br>[0.009; 0.043]- |
| $\beta$ | The $\beta$ parameter of the beta distribution that determines $s$ for P-elements outside piRNA clusters | 10.743<br>[10.128; 15.860] | 17.379<br>[10.026; 19.862] |
| $\vert \bar{s}\vert $ | Mean selection coefficient for new P-element insertions outside piRNA clusters; ($E(\vert s\vert ) = \vert \bar{s}\vert = \alpha/(\alpha + \beta)$) | 0.041<br>[0.030; 0.046] | 0.002<br>[0.001; 0.002] |
| 95<sup>th</sup> percentile $\vert s\vert $ | 95<sup>th</sup> percentile of $\vert s\vert $ for new P-element insertions outside piRNA clusters; | 0.160<br>[0.115; 0.173] | 0.011<br>[0.000; 0.011] |
| Neutral fraction | Expected fraction of effectively neutral ($\vert s\vert \times N_e \leq 1$) P-element insertions outside of piRNA clusters | 0.199<br>[0.170; 0.310] | 0.898<br>[0.892; 0.971] |
| $\vert \bar{s}_{\text{effective}}\vert $ | Mean selection coefficient for new P-element insertions outside piRNA clusters where selection is strong enough to overcome genetic drift ($\vert s\vert \times N_e > 1$) | 0.051<br>[0.038; 0.055] | 0.021<br>[0.019; 0.031] |
| NRMSE<sub>sum</sub> | Fit metric: NRMSE<sub>sum</sub> = NRMSE 1<sup>st</sup> + NRMSE 2<sup>nd</sup> wave<br>NRMSE = normalized root mean square error | 1.340<br>[1.340; 1.509] | 2.313<br>[2.313; 2.388] |
| Best NRMSE<sub>sum</sub> Rank | Rank of the parameter combination with the minimum NRMSE<sub>sum</sub> within group | 1<br>[1; 100] | 38394<br>[38394; 56323] |

**Table S4.** Summary of top-ranked parameter combinations from the parameter exploration with the Gaussian Process model trained on the extended individual-based P-element invasion model that includes additional parameters for excision probability (*v*) and dominance (*h*). Parameter combinations are grouped *post hoc* by mean selection efficacy (2 × *h* × |*s̅*| × *N*_*e*_) under the assumption of *N_e_* = 442. Brackets indicate the range across the top 100 parameter combinations ranked by lowest NRMSE_sum_. Variation among the top 100 parameter combinations reflects that agreement between GP predictions and experimental data depends on specific combinations of model parameters, and that the fitness effects of new P-element insertions arise from the product of *h* × *s*, limiting the interpretability of *s* or *h* individually.

| Parameter/<br>Metric | Description | Mean selection efficacy |  |
| --- | --- | --- | --- |
| | | $2 \times h \times \vert \bar{s}\vert \times N_e > 1$ | $2 \times h \times \vert \bar{s}\vert \times N_e \leq 1$ |
| $p_{\text{carrier}}$ | The probability that an isofemale line carries the P-element at the beginning of the simulation | 0.215<br>[0.153; 0.260] | 0.172<br>[0.150; 0.335] |
| $f_{\text{regulatory}}$ | The piRNA cluster size | 0.014<br>[0.010; 0.022] | 0.087<br>[0.076; 0.100] |
| $u$ | Transition probability of a single P-element per generation | 0.339<br>[0.268; 0.391] | 0.352<br>[0.308; 0.500] |
| $v$ | Excision probability at the original insertion site given that a P-element transposes | 0.188<br>[0.004; 0.244] | 0.071<br>[0.002; 0.241] |
| $\alpha$ | The $\alpha$ parameter of the beta distribution that determines the selection coefficient $s$ for P-elements outside piRNA clusters | 0.323<br>[0.284; 0.499] | 0.429<br>[0.002; 0.499] |
| $\beta$ | The $\beta$ parameter of the beta distribution that determines $s$ for P-elements outside piRNA clusters | 11.766<br>[10.072; 19.976] | 11.058<br>[10.023; 19.285] |
| $h$ | Dominance coefficient | 0.749<br>[0.488; 0.967] | 0.019<br>[0.002; 0.978] |
| $\vert \bar{s}\vert $ | Mean selection coefficient for new P-element insertions outside piRNA clusters; ($E(\vert s\vert ) = \vert \bar{s}\vert = \alpha/(\alpha + \beta)$) | 0.027<br>[0.018; 0.044] | 0.037<br>[0.000; 0.045] |
| 95<sup>th</sup> percentile $\vert s\vert $ | 95<sup>th</sup> percentile of $\vert s\vert $ for new P-element insertions outside piRNA clusters; | 0.119<br>[0.079; 0.165] | 0.149<br>[0.000; 0.171] |
| Neutral fraction | Expected fraction of effectively neutral ($2 \times h \times \vert s\vert \times N_e \leq 1$) P-element insertions outside of piRNA clusters | 0.300<br>[0.168; 0.334] | 0.786<br>[0.681; 1.000] |
| $\vert \bar{s}_{\text{effective}}\vert $ | Mean selection coefficient for new P-element insertions outside piRNA clusters where selection is strong enough to overcome genetic drift ($2 \times h \times \vert s\vert \times N_e > 1$) | 0.038<br>[0.026; 0.053] | 0.120<br>[0.019; 0.509] |
| NRMSE<sub>sum</sub> | Fit metric: NRMSE<sub>sum</sub> = NRMSE 1<sup>st</sup> + NRMSE 2<sup>nd</sup> wave<br>NRMSE = normalized root mean square error | 1.353<br>[1.353; 1.418] | 2.295<br>[2.295; 2.338] |
| Best NRMSE<sub>sum</sub> Rank | Rank of the parameter combination with the minimum NRMSE<sub>sum</sub> within group | 1<br>[1; 100] | 205378<br>[205378; 258411] |

### Supplementary Figures

**Figure S1.**
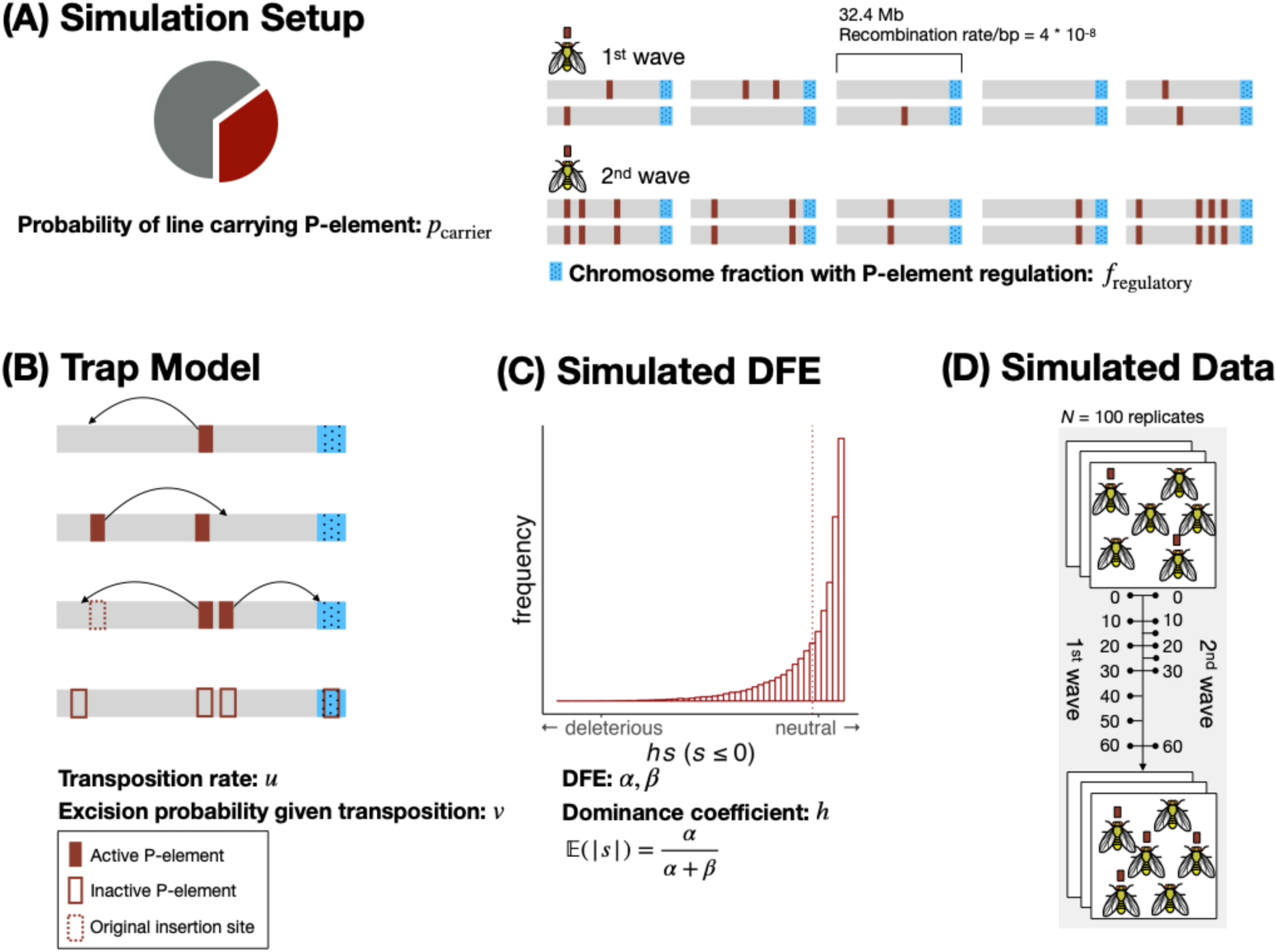
Schematic overview of the extended individual-based simulation model mimicking the P-element invasion dynamics in our Experimental Evolution experiments, including additional model parameters for excision probability (*v*) and dominance (*h*). **(A) Simulation Setup:** Ancestral outbred populations are generated by mixing 200 isofemale lines (five flies each). Each line carries the P-element with probability *p*_carrier_. We modeled diploid individuals with five chromosomes, each with a fixed length of 32.4 Mb and a recombination rate of 4×10⁻⁸ per bp per generation. For the 1^st^ wave, P-element insertions are assumed to be heterozygous. For the 2^nd^ wave, since the experiment started with isofemale lines that had been maintained at small populations sizes for 4.5 years [11], the model assumes that all P-element insertions are homozygous due to increased inbreeding and the likely establishment of a defense mechanism. The parameter *f*_regulatory_ defines the fraction of each chromosome with P-element-regulatory properties (piRNA clusters; blue rectangles). **(B) Trap Model:** The P-element remains active (filled rectangle) unless one of its copies transposes into a piRNA cluster (blue rectangles). The probability that a single P-element undergoes transposition in a given generation is controlled by the transposition rate *u*. If transposition occurs, the probability of a clean excision event at the original insertion site — represented by an unfilled rectangle with dotted red borders — is determined by the parameter *v*. Once a piRNA cluster acquires a single P-element insertion, all P-elements in the genome are immediately inactivated (unfilled rectangles with solid red borders). **(C) Simulated Distribution of Fitness Effects (DFE):** The DFE for new P-element insertions is modeled using a beta distribution with shape parameters *α* and *β,* which define the selection coefficient *s*. Each new P-element insertion is also assigned a dominance coefficient *h* ranging between 0 and 1. Positive selection is not considered in our model. **(D) Simulated Data:** The simulation output used in the analyses contains the average P-element copy number per haploid genome across 100 simulation runs, taken at the same time points used in the two Experimental Evolution studies.

**Figure S2.**
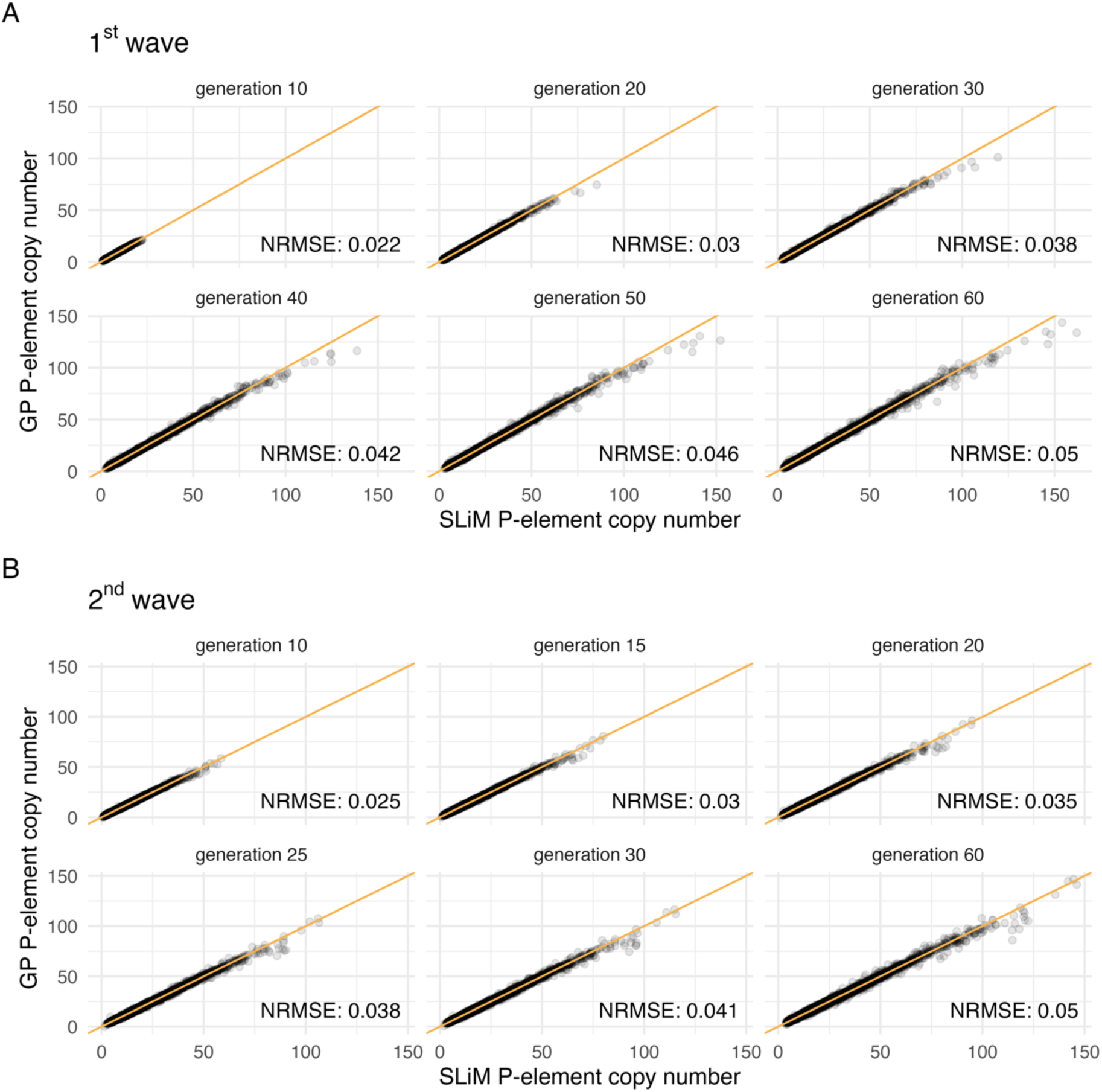
Gaussian Process (GP) performance: A test dataset consisting of 10000 data points was simulated with the extended individual-based model (*x* axis) for the (A) 1^st^ wave and (B) 2^nd^ wave and compared to the predictions of the GP (*y* axis). One data point in this test dataset comprises a specific combination of seven model parameters (Table S1) and the corresponding predicted P-element copy numbers for six time points. The observed and predicted P-element copy numbers are shown (gray dots) for each of those six time points (the six panels), with amber lines indicating the identity line (*x = y*). Normalized root mean square errors (NRMSE) for the time points are shown at the bottom right of the corresponding panels. The analysis shows that the GP can predict the copy number observed in the individual-based model very accurately. Only at extremely high copy numbers — well beyond the empirical estimates (Figure 2) — can a slight underestimation be observed.

**Figure S3.**
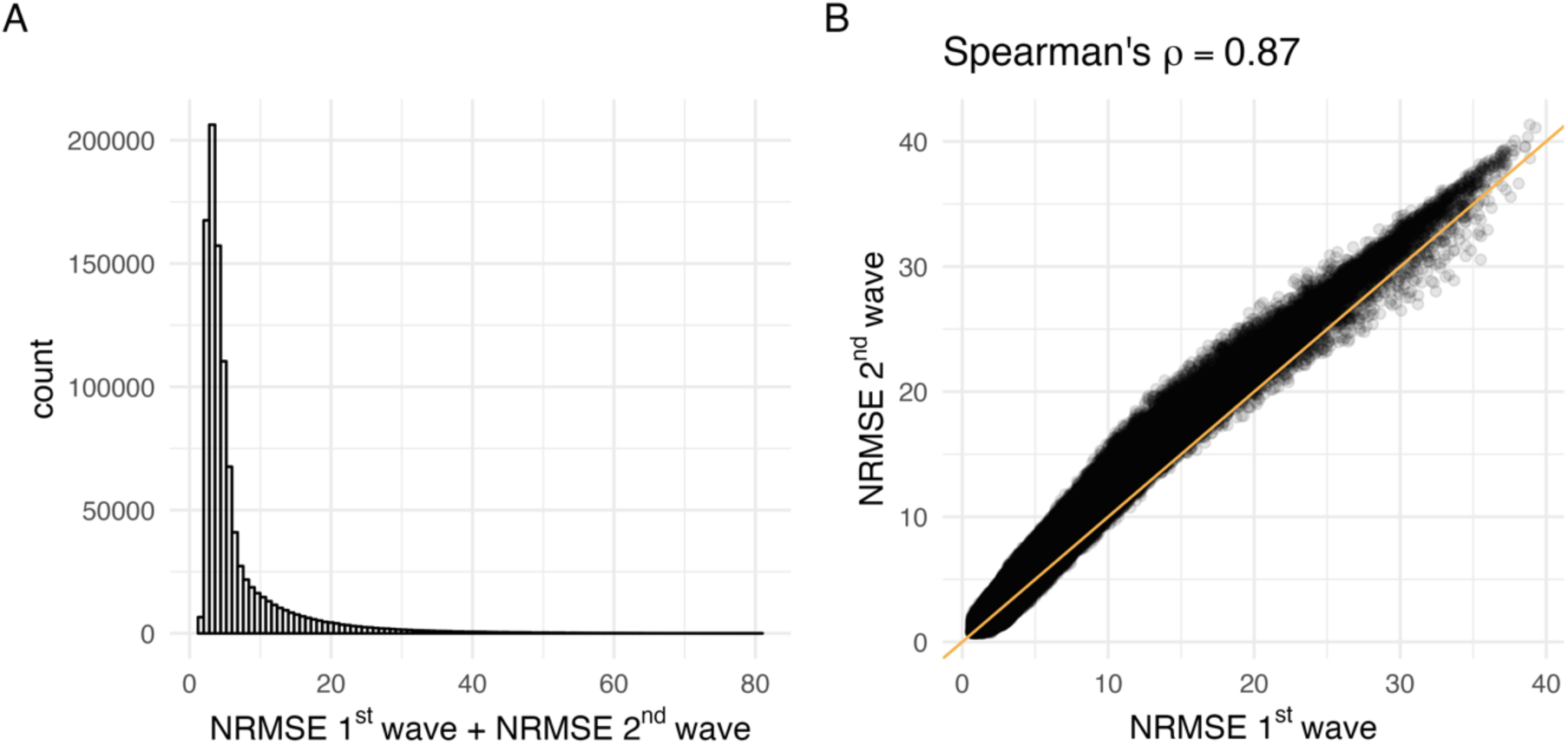
Normalized root mean square error (NRMSE) for GP predictions using 10⁶ different parameter combinations. (A) Distribution of NRMSE sums, showing variation in the agreement between GP predictions and empirical data. While most values are moderate, a few combinations have very low or very high NRMSE values. (B) Comparison of NRMSE for the 1^st^ wave (*x* axis) versus the 2^nd^ wave (*y* axis). Each point represents the NRMSEs for one of the 10⁶ data points. The amber line represents the identity line (*x = y*). Overall, there is strong agreement between the NRMSEs, with a Spearman rank correlation coefficient of 0.87.

**Figure S4.**
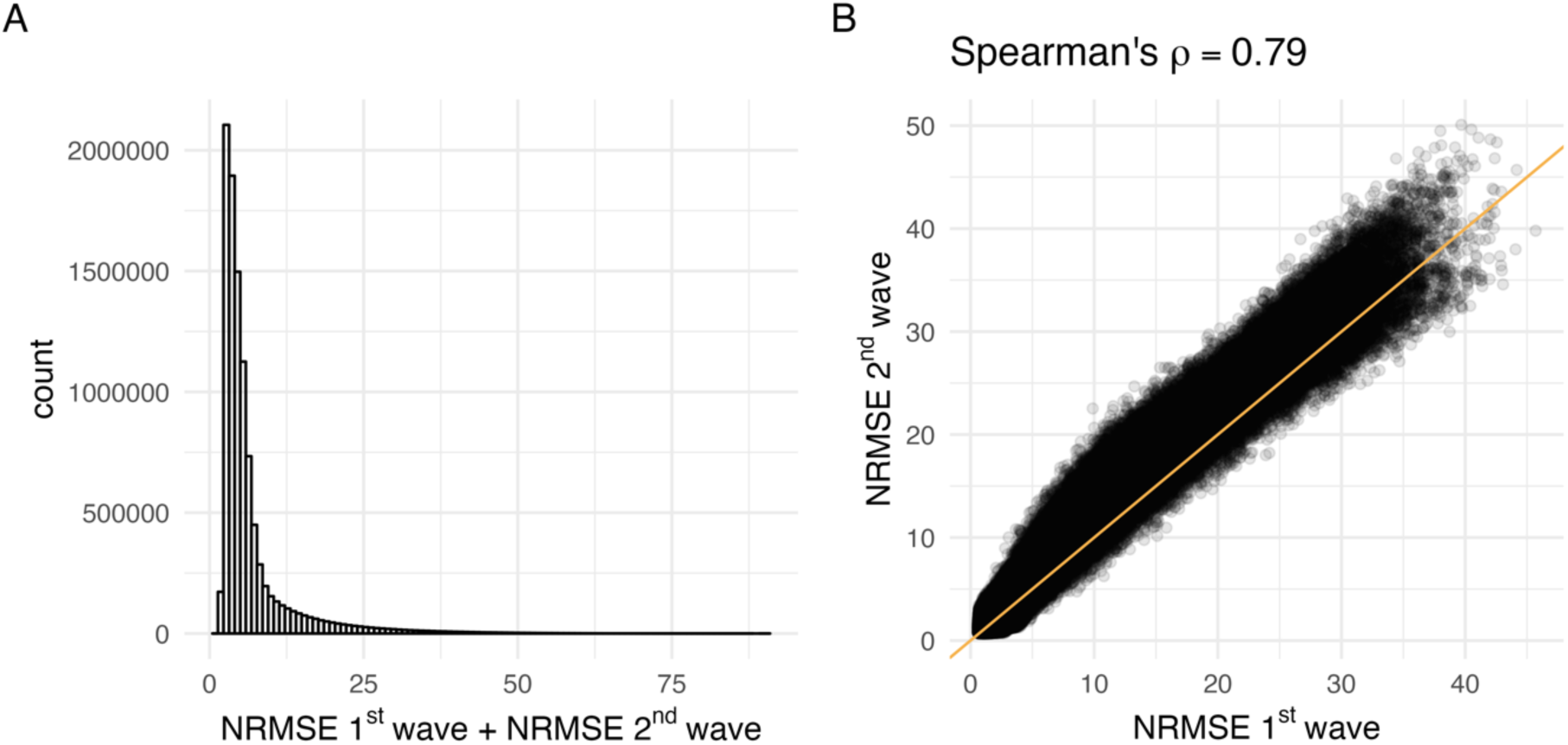
Normalized root mean square error (NRMSE) for predictions made by GPs trained on simulation outcomes from the extended individual-based model. (A) Distribution of NRMSE sums, showing variation in the agreement between GP predictions using 10^7^ different parameter combinations and empirical data. While most values are moderate, a few combinations have very low or very high NRMSE values. (B) Comparison of NRMSE for the 1^st^ wave (*x* axis) versus the 2^nd^ wave (*y* axis). Each point represents the NRMSEs for one of the 10^7^ data points. The amber line represents the identity line (*x = y*). Overall, there is strong agreement between the NRMSEs, with a Spearman rank correlation coefficient of 0.79.

**Figure S5.**
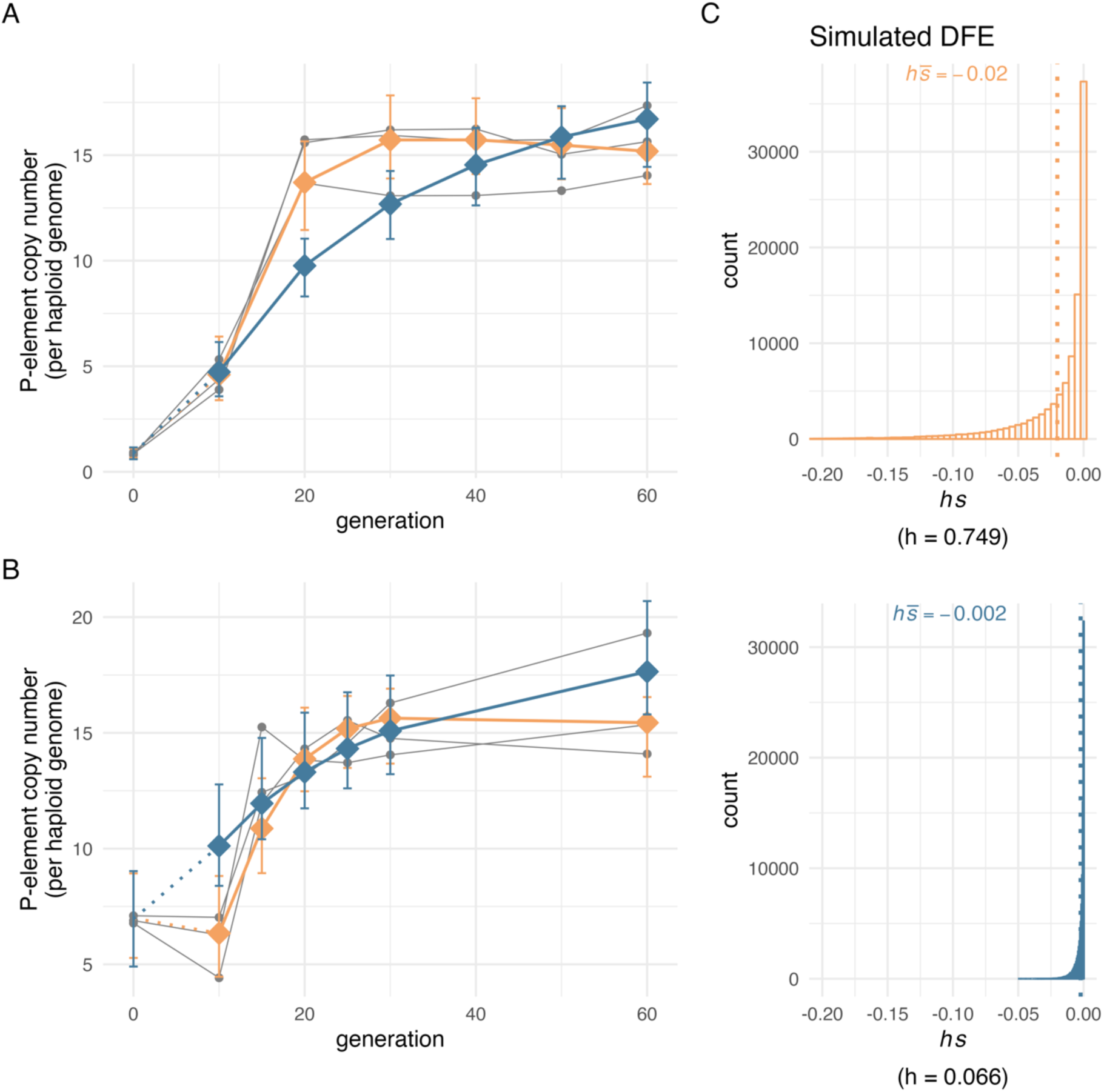
Gaussian Process (GP) predictions provide a good fit to the empirical data — the top-ranked parameters were identified using 1^st^ and 2^nd^ wave data. GPs were trained on simulation outcomes of the extended individual-based model. GP predictions from the parameter combinations with the lowest NRMSE_sum_ when compared against empirical data are shown for scenarios with expected effective purifying selection (2 × *h* × |*s̅*| × *N*_*e*_ > 1, amber), and for scenarios without expected effective purifying selection (2 × *h* × |*s̅*| × *N*_*e*_ ≤ 1, steel blue) assuming *N_e_* = 221 [54]. Each grey line represents an empirical evolution replicate, with sequenced time points indicated by dots. GP predictions are indicated by connected colored diamonds. The error bars show the range between the 2.5^th^ and 97.5^th^ percentiles of P-element copy number trajectories simulated with the individual-based model. These simulations were run with the same parameters as those used for GP prediction. The error bars are shown at the same time points where empirical data were sequenced. Note that GP predictions do not cover generation 0; instead, predictions from generation 10 are connected to the individual-based simulation averages at generation 0 to aid visual interpretation. (A) 1^st^ wave experiment. (B) 2^nd^ wave experiment. (C) Simulated distribution of fitness effects (DFE) used in (A) and (B). Due to selection, the actual distribution of selection coefficients for segregating P-element insertions in the simulated populations will be skewed toward 0. Results are robust across the 100 parameter combinations with the lowest NRMSE sums (Figure S7).

**Figure S6.**
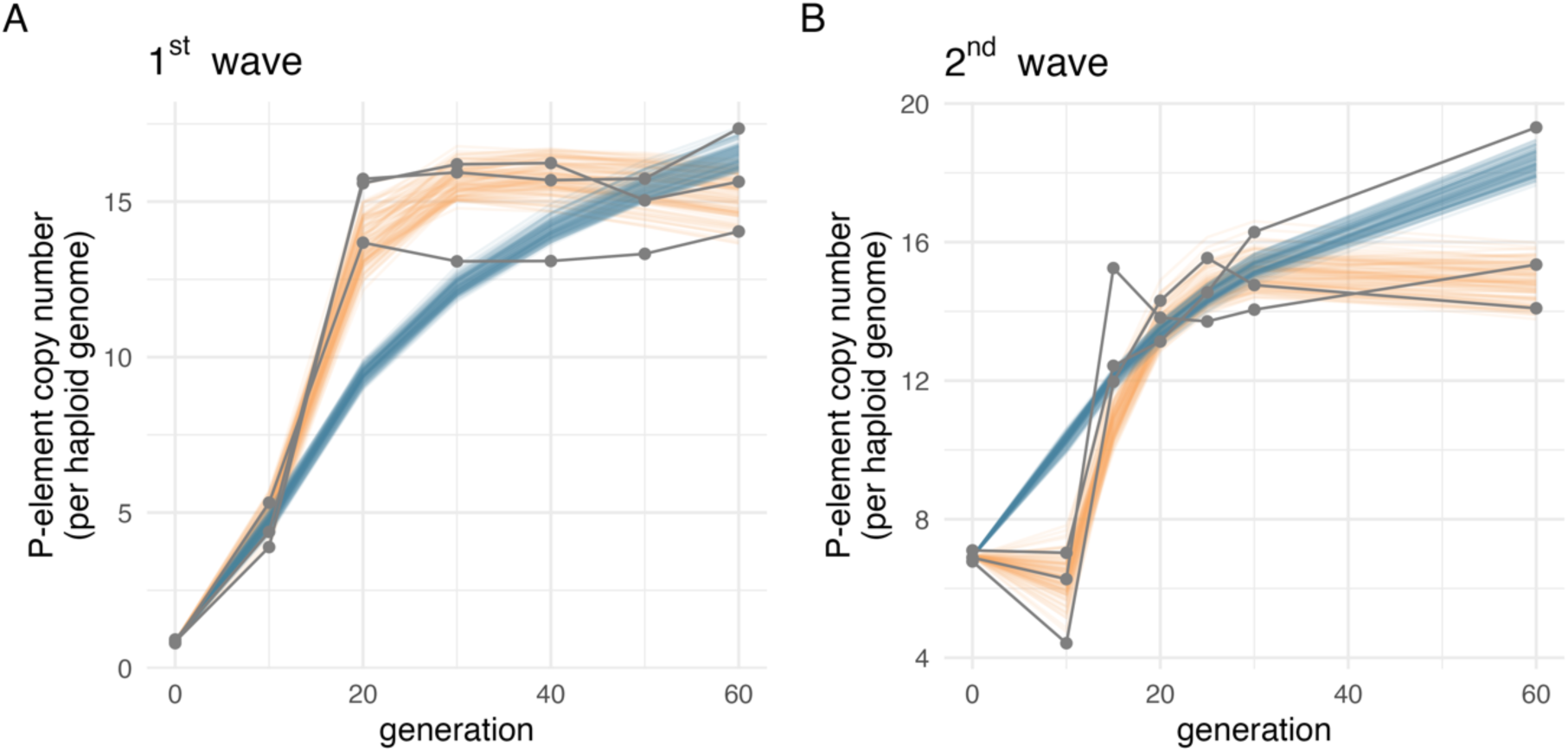
Individual Gaussian Process (GP) predictions for the top-ranked parameter combinations identified through parameter exploration. Predictions are grouped *post hoc* based on the mean selection efficacy: scenarios with expected effective purifying selection (|*s̅*| × *N*_*e*_ > 1, amber) and scenarios without (|*s̅*| × *N*_*e*_ ≤ 1, steel blue), assuming *N*_e_ = 221 [54]. For each group, predictions from the 100 top-ranked parameter combinations (based on lowest NRMSE_sum_, Table 2) are shown. Each grey line represents an empirical evolution replicate, with sequenced time points indicated by dots. Note that GP predictions do not cover generation 0; instead, predictions from generation 10 are connected to the empirical average at generation 0 to aid visual interpretation.

**Figure S7.**
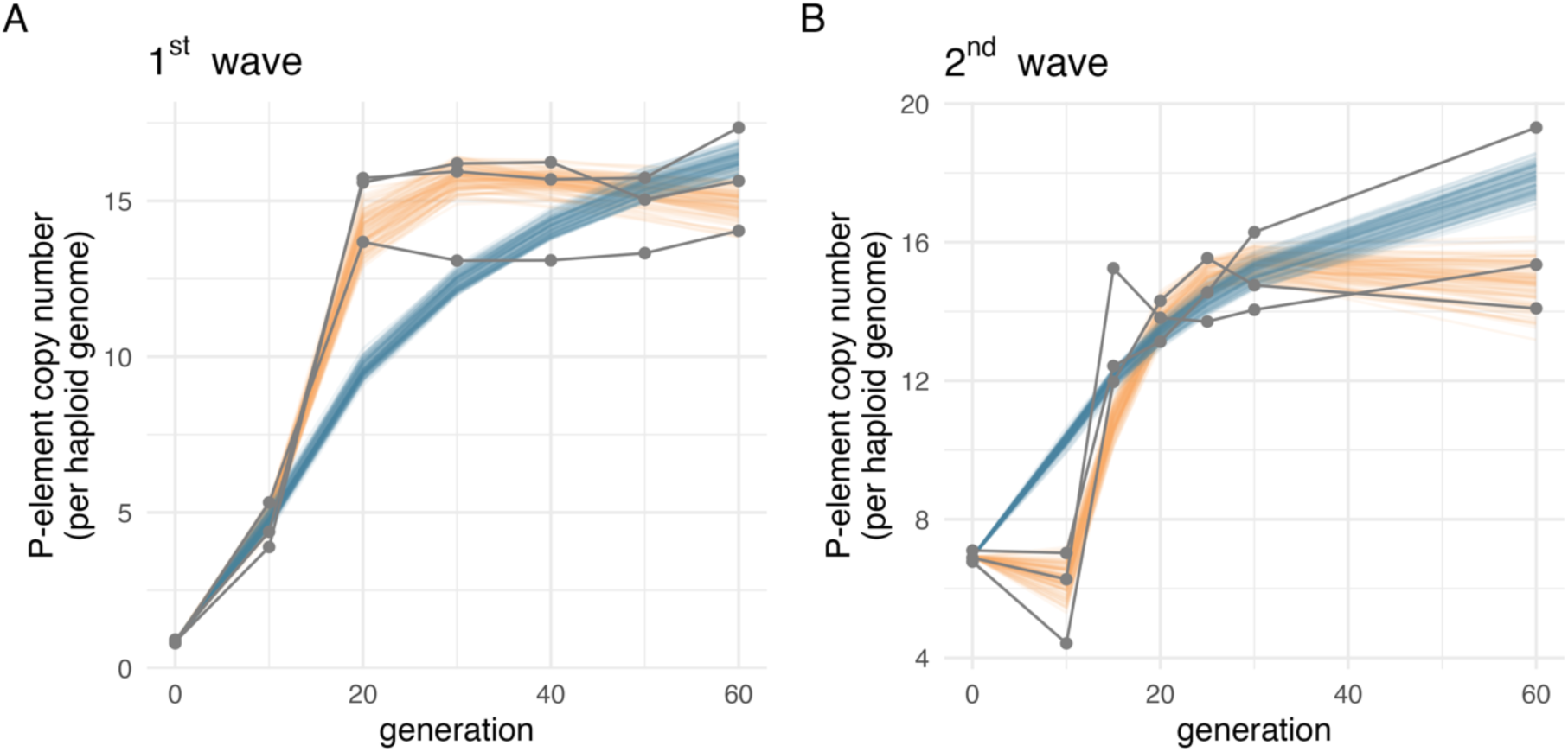
Individual Gaussian Process (GP) predictions for the top-ranked parameter combinations identified through parameter exploration. GPs were trained on simulation outcomes of the extended individual-based model. Predictions are grouped *post hoc* based on the mean selection efficacy: scenarios with expected effective purifying selection (2 × *h* × |*s̅*| × *N*_*e*_ > 1, amber) and scenarios without (2 × *h* × |*s̅*| × *N*_*e*_ ≤ 1, steel blue), assuming *N*_e_ = 221 [54]. For each group, predictions from the 100 top-ranked parameter combinations (based on lowest NRMSE_sum_, Table S2) are shown. Each grey line represents an empirical evolution replicate, with sequenced time points indicated by dots. Note that GP predictions do not cover generation 0; instead, predictions from generation 10 are connected to the empirical average at generation 0 to aid visual interpretation.

